# Cytoplasmic roles of HSATIII RNAs in RNA granule assembly and production of actin cytoskeleton-associated repeat-containing proteins

**DOI:** 10.1101/2025.11.12.688122

**Authors:** Kensuke Ninomiya, Shungo Adachi, Toru Suzuki, Sylvie Souquere, Gerard Pierron, Toshifumi Inada, Tetsuro Hirose

## Abstract

HSATIII architectural lncRNAs are nuclear short-tandem repeat ncRNAs induced upon thermal stress and serve as scaffolding molecules for nuclear stress bodies (nSBs), stress-induced membraneless organelles in the nucleus. We report that HSATIII lncRNAs are exported to the cytoplasm during stress recovery, where they form novel cytoplasmic bodies with nucleocytoplasmic RNA-binding proteins, including PURA, PURB, and m^6^A reader proteins. Recruitment of ALYREF to nSBs promotes HSATIII RNA export. These HSATIII-based granules, named HERALDs (Heat-removal-associated scaffolds), are transported along microtubules and accumulate at distal cellular regions. We further show that cytoplasmic HSATIII RNAs are translated into MEWNG-rich proteins, which interact with actin filaments and form complexes with HSPA8 and stress granule proteins such as ZC3HAV1 and ATXN2L. Together, these findings reveal previously unrecognized cytoplasmic roles of HSATIII RNAs, functioning both as lncRNAs to scaffold RNA granules and as mRNAs to produce proteins during the late phase of thermal stress recovery.

## INTRODUCTION

HSATIII long noncoding RNAs (lncRNAs) are primate specific transcripts produced from the pericentromeric satellite III regions of several chromosomes under various stress conditions^1–4^. The majority of HSATIII lncRNA sequence consists of GGAAU repeats that are interspersed with less frequent CAACCCGARU repeats^5^. Transcribed HSATIII lncRNAs remain stable in the nucleus and recruit specific RNA-binding proteins (RBPs) including SAFB, SRSFs and HNRNPs, thereby acting as architectural RNAs^6^ that assemble membraneless organelles called nuclear stress bodies (nSBs)^7–9^. The protein components of nSBs, identified by Chromatin Isolation by RNA purification (ChIRP) of HSATIII architectural lncRNA followed by mass-spectrometry (HSATIII-ChIRP-MS), appeared to change dynamically during the temperature shift from thermal stress to recovery^10,11^.

Our transcriptome analysis of cells in which HSATIII was knocked down, and therefore unable to form nSBs, revealed that nSBs repress splicing of specific introns in hundreds of pre-mRNAs, predominantly during the early recovery phase after the thermal stress removal^11^. As underlying mechanisms, two biochemical modifications occur within nSBs specifically during the post-stress recovery phase^10,11^.

As the first mechanism, nSBs concentrate specific sets of SR splicing factors (SRSFs) and SR-related proteins under thermal stress conditions and recruit CLK1 kinase specifically in the post-stress recovery phase to rapidly rephosphorylate SRSFs especially SRSF9, which were dephosphorylated upon thermal stress^11^. In turn, the rephosphorylated SRSF9 represses splicing of more than one hundred of pre-mRNAs. As the second mechanism, N6-methyladenoosine (m^6^A) writer complex is recruited into nSBs and adenosines in GGAAU repeat of HSATIII lncRNAs are partially m^6^A-modified especially in the post-stress recovery phase, Thereby, nSBs sequestrate nuclear m^6^A reader protein YTHDC1, resulting in repressing YTHDC1-dependent splicing of pre-mRNAs^10,12^. The role of nSBs in regulating pre-mRNA splicing is most evident during the early recovery phase, 1 hour after stress removal, when they are readily detectable in the nucleus. HSATIII is also reported to be involved in regulation of transcription, other RNA processing events and chromatin segregation probably through distinct mechanisms^13–15^, suggesting that it has multifaceted molecular functions in the nucleus.

Here, we report that a population of HSATIII RNAs is exported to the cytoplasm during the extended recovery period following thermal stress removal, where they assemble novel cytoplasmic RNP granules together with nucleo/cytoplasmic RBPs such as PURA, PURB, FXR1/2 and cytoplasmic m^6^A readers, YTHDFs. These cytoplasmic HSATIII granules, termed HERALD bodies, are typically detected 4 hours after stress removal at the specific sites along the distal edge of the cells. Furthermore, we also demonstrated that cytoplasmic HSATIII RNAs are likely translated into MEWNG repeat-containing polypeptides (HSATIII proteins). Thus, these findings reveal previously unrecognized functions of HSATIII RNA in the cytoplasm, acting as protein-coding mRNAs in addition to their roles as lncRNAs that assemble RNA granules in both the nucleus and cytoplasm.

## RESULTS

### Emergence of novel cytoplasmic RNP granules assembled on HSATIII lncRNAs during extended recovery from thermal stress

HSATIII lncRNAs are transcribed under thermal stress conditions and remain stable in the nucleus as nSBs for several hours after the stress removal^16^. We found that HSATIII RNA foci were also detectable in the cytoplasm around 4 hours after the stress removal, especially at the distal ends (Figure 1A, arrowheads). Since cytoplasmic HSATIII RNA foci were frequently observed at the adhesive surface between the cells, it was difficult to count the number of the cytoplasmic foci of each cell. Alternatively, we estimated the number of the cytoplasmic foci per the number of the cells from at least three image fields and also quantified their sizes. The number and size of cytoplasmic HSATIII RNA foci were increased from 2 hours after the stress removal (Figure 1A and 1B). We previously reported HSATIII-ChIRP-MS data at various time points of thermal stress and recovery, and identified nucleocytoplasmic RBPs such as PURA, PURB and TDP-43 that were detected in the ChIRP fraction preferentially in the recovery phase after stress removal^10^. We confirmed by western blot of the HSATIII-ChIRP samples that the amounts of PURA, PURB and TDP-43 in the HSATIII-ChIRP fractions were increased at the 4-hour recovery phase in compared to the 1-hour recovery phase after thermal stress (Figure 1C), suggesting the possibility that HSATIII lncRNAs form the cytoplasmic foci together with these cytoplasmic RBPs. Then, we checked the colocalizations of these RBPs with HSATIII lncRNAs in HeLa cells the 4-hour recovery phase. PURA and PURB were almost completely colocalized with cytoplasmic HSATIII RNA foci, but barely with HSATIIII RNAs in the nucleus (Figure 2A). On the other hand, TDP-43 was partly colocalized with HSATIII lncRNAs both in the nucleus and cytoplasm (Figure 2B). In the cells continuously exposed to 6-hour thermal stress without recovery, neither cytoplasmic HSATIII RNA nor PURA foci were detected (Figure S1A), suggesting that the recovery process at 37℃ is required for the formation of these cytoplasmic foci. These foci remained stable at least 8 hours after the stress removal but almost disappeared in 24 hours (Figure S1B).

**Figure 1.**
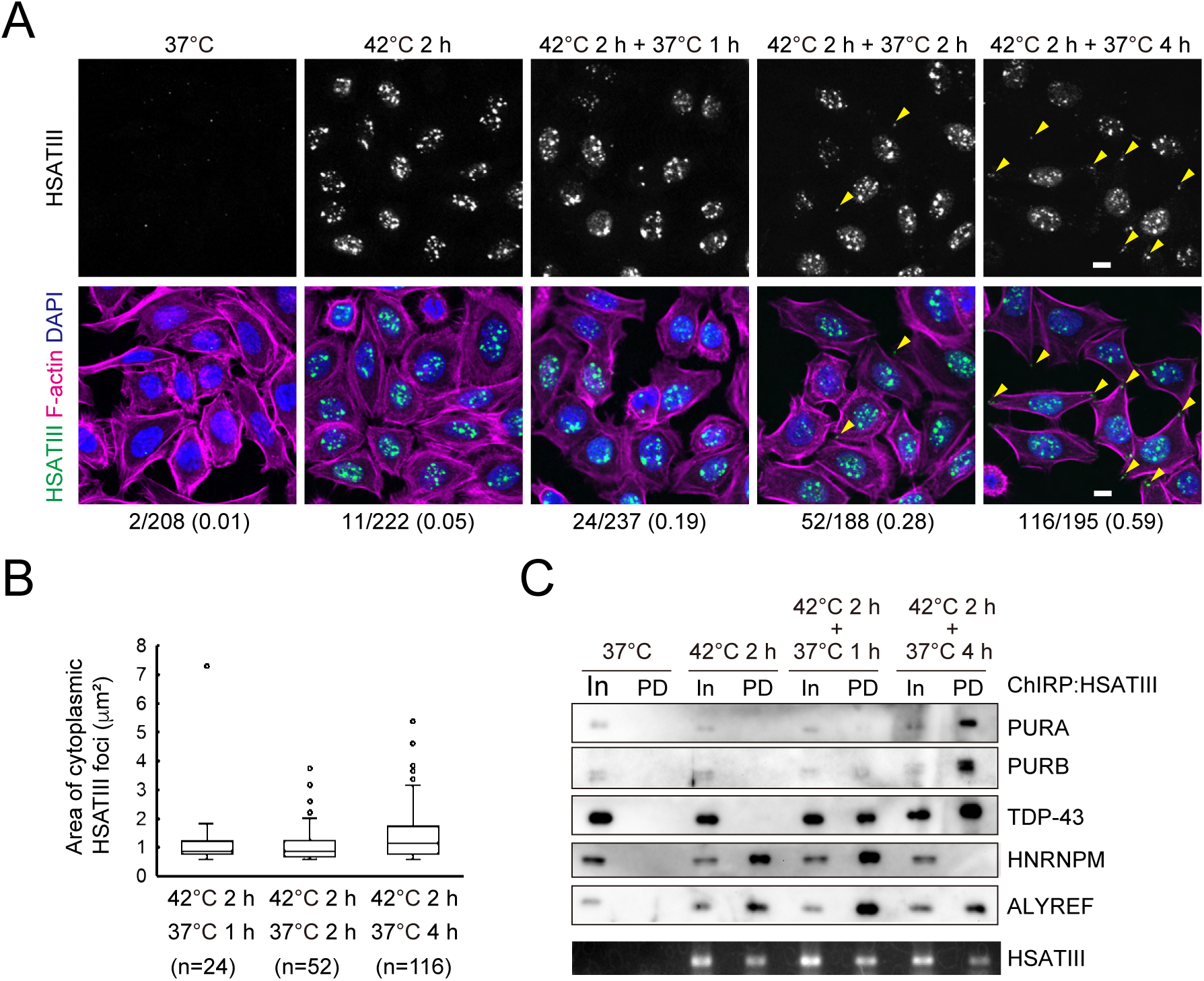
Formation of cytoplasmic HSATIII RNA foci in thermal stress recovery. (A) Subcellular distribution of HSATIII RNAs. HeLa cells were exposed to thermal stress (42 °C for 2 h followed by recovery at 37 °C). HSATIII RNAs were stained by FISH. F-actin and nuclei were stained by phalloidin rhodamine and DAPI, respectively. The total number of the cytoplasmic HSATIII RNA foci per that of nuclei in the obtained microscopic images is indicated. Scale bar: 10 mm (B) Box plot of area of individual cytoplasmic HSATIII RNA foci (0.5 - 10mm^2^) in (B). The numbers of the quantified foci (n) are indicated below. (C) ChIRP-western blot. HSATIII-ChIRP was performed from HeLa cells prepared at various time points of thermal stress and recovery as indicated. Western blot was performed using a specific antibody as indicated. Semi-quantitative RT-PCR was performed to detect HSATIII RNAs (bottom panel). PD: pulldown. Input (In) :1 % for WB, 100% for RT-PCR

**Figure 2.**
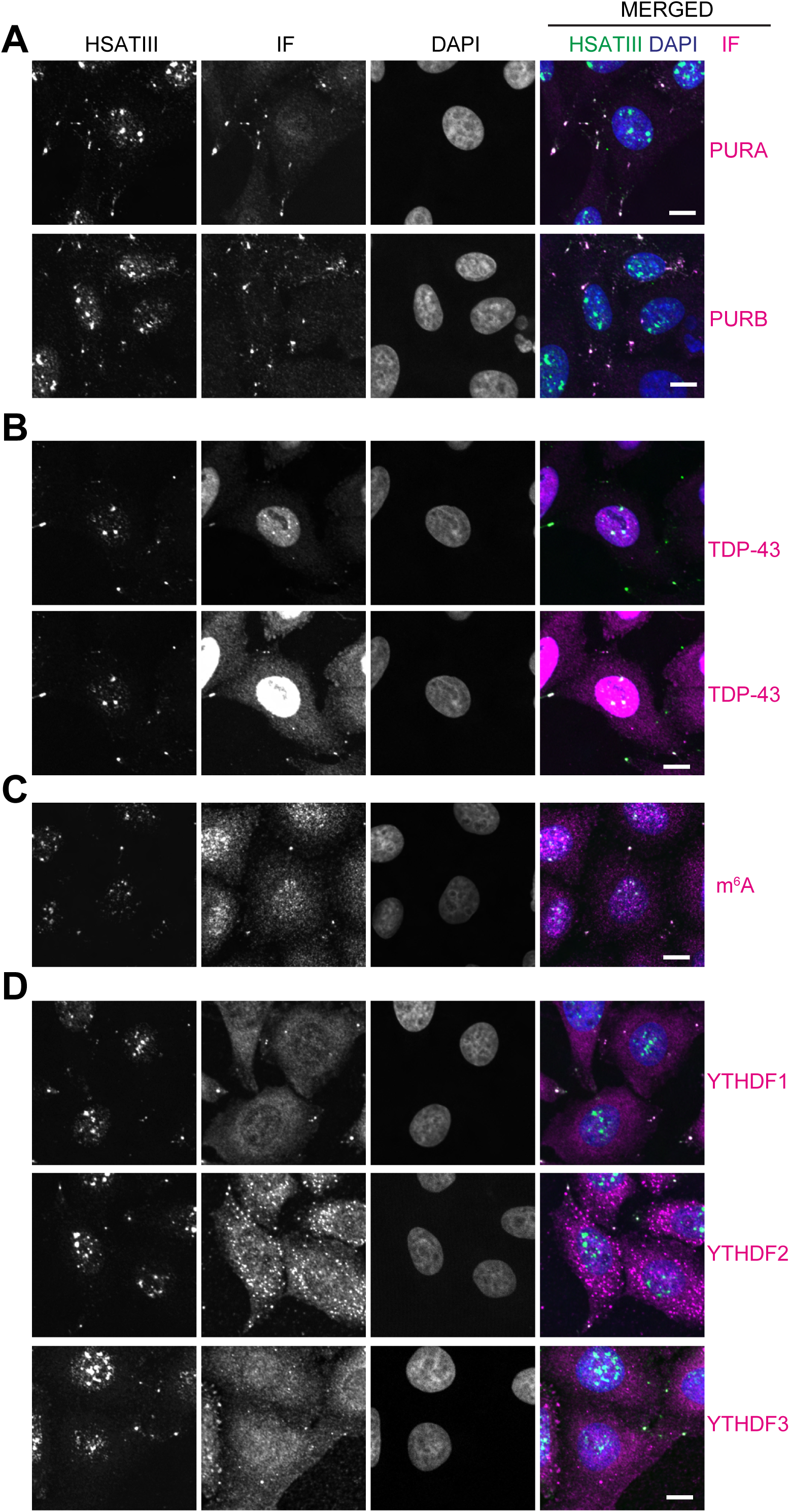
RBPs and nucleotide colocalized with cytoplasmic HSATIII lncRNA foci. (A-D) HeLa cells (cultured at 42 °C for 2 h followed by recovery at 37 °C for 4 h) were stained by HSATIII-FISH and IF using a specific antibody against each protein and m^6^A nucleotide. Nuclei were stained by DAPI. Scale bar: 10 mm

We previously reported that HSATIII lncRNAs are methylated by m^6^A writer complex and recruit the nuclear m^6^A reader YTHDC1 into nSBs^10^. m^6^A immuno-positive foci were detected in the cytoplasmic HSATIII RNA foci as well as nSBs (Figure 2C). Cytoplasmic m^6^A reader proteins (YTHDF1, YTHDF2 and YTHDF3) also colocalized with cytoplasmic HSATIII lncRNA foci, whereas YTHDC1 did not (Figure 2D and Figure S2A). This suggests that cytoplasmic HSATIII lncRNAs, and/or some colocalizing RNAs, are m^6^A-modified and recognized by cytoplasmic m^6^A reader proteins, which may replace YTHDC1. Additionally, we confirmed that previously-reported major nSB components such as SAFB, SRSF1, SRSF9, and ALYREF were colocalized with HSATIII lncRNAs only in the nucleus but not in the cytoplasm (Figure S2B-S2E). HNRNPM, another major component of nSBs, was diffused in the nucleus 4 hrs after the stress removal and barely colocalized with the nuclear or cytoplasmic HSATIII RNA foci (Figure S2F), consistently with the result of ChIRP-WB (Figure 1C). These observations suggest that the protein components of cytoplasmic HSATIII RNA granules differ substantially from those of nSBs, with some exceptions such as TDP-43.

### HSATIII RNAs and PURA/B form a novel cytoplasmic body, HERALD

HSATIII lncRNAs act as architectural RNAs that are essential for assembling nSBs^6^. To determine whether HSATIII RNAs also play architectural roles in the assembly of cytoplasmic HSATIII RNA foci, we examined the effect of HSATIII KD using antisense oligonucleotides on PURA foci at the 4-hour recovery phase after stress removal. The number of cytoplasmic PURA foci was markedly reduced in HSATIII KD cells (Figure 3A, arrowheads), suggesting that HSATIII lncRNAs also act as architectural RNA for cytoplasmic foci, as well as for nSBs. Next, we tested whether PURA and PURB are also required for the formation of the cytoplasmic HSATIII RNA granules. Considering that PURA and PURB are paralogues and may be functionally redundant, we performed double KD of PURA and PURB using siRNAs (siPURA/B) and analyzed the effect on HSATIII RNA granule formation. As a result, PURA/B double KD decreased the number and size of the cytoplasmic HSATIII RNA foci without affecting nSB assembly (Figure 3B and 3C). These suggest that HSATIII lncRNAs and PURA/B cooperatively contribute to the RNP granule formation in the cytoplasm.

**Figure 3.**
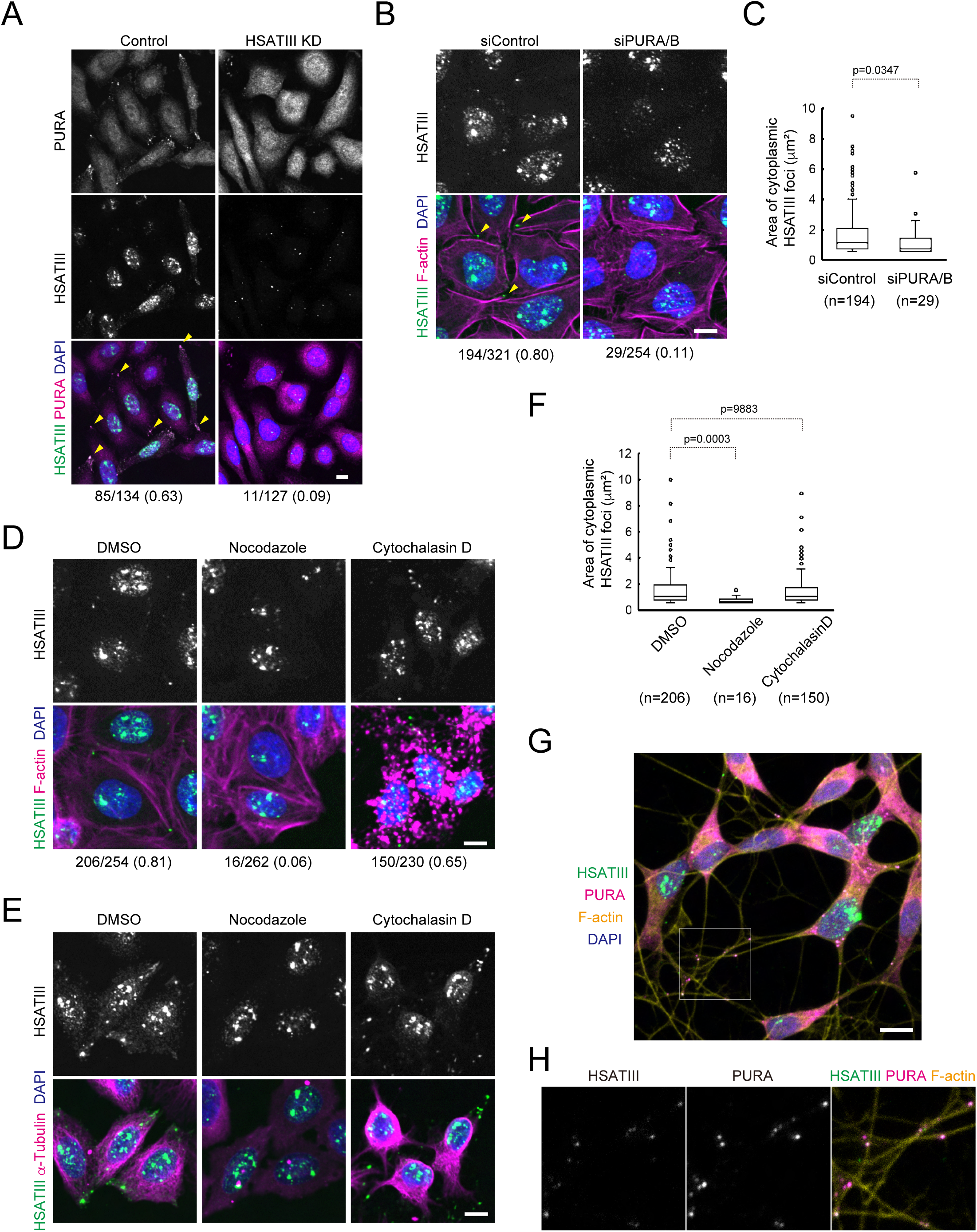
Essential factors for formation of cytoplasmic HSATIII RNP foci. (A) HSATIII-dependent formation of the cytoplasmic PURA foci. HeLa cells were transfected with HSATIII ASO or control sense oligonucleotide, and cultured for 16 h, exposed to thermal stress (42 °C for 2 h, followed by recovery at 37 °C for 4 h), and stained by HSATIII-FISH and IF using PURA antibody. Nuclei were stained by DAPI. Scale bar: 10 □m. The number of cytoplasmic foci per that of nuclei obtained from at least three microscopic images is indicated (the same hereinafter). (B) The effect of PURA/B double knockdown on cytoplasmic HSATIII RNP foci. HeLa cells were transfected with siRNAs against PURA and PURB (10 nM for each) or control siRNA (20 nM), and cultured for 48 h, exposed to thermal stress (42 °C for 2 h, followed by recovery at 37 °C for 4 h), and stained as described in Figure 1A. Scale bar: 10 □m. (C) Box plot of area of individual cytoplasmic HSATIII foci (0.5 - 10 mm^2^) in (B). The numbers of quantified foci are indicated below. p-value (Mann–Whitney U-test) is shown above. (D, E) Microtubule-dependent formation of cytoplasmic HSATIII RNA foci. HeLa cells were exposed to thermal stress (42 °C for 2 h), recovered for 4 h at 37 °C in the presence of DMSO, Nocodazole (10 mg/ml) or Cytochalasin D (2 mg/ml), and stained by HSATIII-FISH and IF using anti-tubulin antibody (E). F-actin (D) and nuclei were stained by phalloidin rhodamine and DAPI, respectively. Scale bar: 10 mm. (F) Box plot of area of individual cytoplasmic HSATIII foci (0.5 - 10 mm^2^). The numbers of quantified foci are indicated below. P-Value (Kruskal-Wallis test, followed by Dunn’s multiple comparison test) was shown above (G, H) Cytoplasmic HSATIII RNP foci in the processes of neuronally differentiated SH-SY5Y cells. SH-SY5Y cells were differentiated by retinoic acid and BDNF (see method), exposed to thermal stress (42 °C for 2 h), recovered for 8 h at 37 °C and stained by HSATIII-FISH and IF using anti-PURA antibody. F-actin and nuclei were stained by phalloidin iFluor^TM^647 and DAPI, respectively. Scale bar: 10 mm.

PURA/B and TDP43 are also known as components of other stress-responsible membraneless organelles, SG and p-body^17–19^. Hence, we tested whether cytoplasmic HSATIII RNA foci were colocalized with G3BP and DCP1a, the well-known maker proteins of stress granules and p-body, respectively. Under thermal stress conditions, G3BP-positive foci distributed proximally around the nucleus and colocalized with PURA (Figure S3A and S3B). At this time point, HSATIII RNAs exclusively localized in the nucleus. At 4-hour stress recovery phase, G3BP has diffused in the cytoplasm and did not colocalized with HSATIII RNA- or PURA-positive foci, which mainly distributed at the distal areas of the cytoplasm. HSATIII RNA foci were not overlapped with p-bodies stained by an DCP1a-antibody (Figure S3C), even though weak signals of PURA were overlapped with p-bodies (Figure S3D). These indicated that cytoplasmic HSATIII RNA foci are distinct from SGs or p-bodies, although they share common components such as PURA/B and/or TDP43. Therefore, we named this novel HSATIII RNA-dependent cytoplasmic body HERALD (Heat-removal associated scaffold).

### HSATIII lncRNAs are transported along microtubules to the distal sites of the cytoplasm

HERALD bodies were frequently observed at the distal ends of the cells (Figure 1A). PURA and PURB are multifunctional nucleocytoplasmic RBPs^20^ and known as adaptor RBPs, that link RNAs with a microtubule-dependent motor protein kinesin in neuronal cells^21^. These findings raised the possibility that cytoplasmic HSATIII RNAs reaccumulate at the distal ends of cells via microtubule-based RNP transport. To test this possibility, we examined the effect of microtubule destabilization on the formation of HERALD bodies. Following stress removal, the number of HERALD foci in the cytoplasm was decreased when the cells were treated with microtubule polymerization inhibitor Nocodazole during the recovery period (Figure 3D and 3E). The size of the remaining HERALD bodies was also smaller compared to the control (Figure 3E and 3F). In contrast, the number and size of HERALD bodies were hardly affected by Cytochalasin D-induced disruption of actin cytoskeleton (Figure 3D-3F). These suggest that HSATIII RNAs move along microtubules and re-accumulate at the distal ends of HeLa cells. On the other hand, Nocodazole did not disrupt pre-formed HERALD bodies (Figure S3E), suggesting that microtubule is required for re-accumulation and de novo assembly of HSATIII RNPs but not for the maintenance of HERALD bodies. The cytoplasmic foci containing HSATIII RNAs and PURA were also detected along the neurites of neuronally differentiated SH-S5Y5 cells that were heat-treated for 2 hours and then allowed to recover for 8 hours (Figure 3G and 3H), suggesting that HERALD bodies are formed not only at the distal ends of HeLa cells but also in the processes of neuronal cells, where microtubule-based RNP transport mechanisms are highly-developed^22,23^,.

### Cytoplasmic HSATIII RNAs are translated into novel MEWNG-rich proteins

HSATIII RNAs have been classified as long non-coding RNAs, primarily because they were long believed to localize exclusively to the nucleus, where they perform distinct functions described above. However, our observation that HSATIII RNAs are also detected in the cytoplasm during recovery from thermal stress raised a possibility that these RNAs also have the potential to be translated as mRNAs. Since HSATIII RNAs are heterogenous repetitive transcripts consisting predominantly of highly-repetitive GGAAU motifs with less frequent CAACCCGARU motifs as minor repeat sequences, their translated products are predicted to be rich in MEWNG-like repeats. These MEWNG-rich polypeptides would arise from the GGAAU pentamer repeats regardless of reading frames while the less abundant CAACCCGARU-derived motifs would produce amino acid sequences depending on reading frame, as shown in Figure 4A. Accordingly, we generated a polyclonal antibody against a MEWNG repeat polypeptide [(MEWNG)X5] and used it for western blotting to test whether cytoplasmic HSATIII RNAs are translated into MEWNG-rich polypeptides during the recovery phase from stress. As a result, broad signals in the 20-40 kDa range were detected in the extract from cells 4 hours into the recovery phase after thermal stress, whereas no such signals were observed under normal conditions or thermal stress condition without recovery (Figure 4B). In addition, these signals disappeared when the cells were treated with translation inhibitor cycloheximide (CHX) during the recovery period (Figure 4B), indicating that MEWNG-rich proteins are produced within 4 h after the thermal stress removal. Moreover, these signals were not detected in HSATIII KD cells (Figure 4C), indicating that these proteins are HSATIII RNA-derived; thus they are referred to as HSATIII proteins. HSATIII proteins were only modestly detected in cells continuously exposed to thermal stress (42℃ 6 h) without recovery (Figure S4A), suggesting that the recovery condition at 37℃ is required for efficient production of HSATIII proteins, in a manner analogous to HERALD formation (Figure S1A). Time-course analysis by western blotting revealed that HSATIII proteins gradually accumulated between 2 and 8 hours after stress removal, but had nearly disappeared by 24 hours after stress removal (Figure S4B). The production of HSATIII proteins as well as HERALD formation, was prevented upon PURA/B dKD (Figure 4E). Supporting the translation of cytoplasmic HSATIII RNA, we detected HSATIII RNAs in polysome fractions purified from cells in the thermal stress-recovery phase, when the production of HSATIII polypeptides was at its peak (42℃ 2 h + 37℃ 2 h) (Figure 4E). Collectively, these results strongly suggest that HSATIII RNAs function as mRNAs and are translated into MEWNG-rich HSATIII proteins during the thermal stress recovery. Importantly, nocodazole treatment, which prevents HERALD formation (Figure 3D-F), did not affect the production of HSATIII proteins (Figure S4C), indicating that HERALD formation is not required for HSATIII RNA translation. On the other hand, we found that FXR1 and FXR2, which are known components of translation-repressed, transportable mRNPs and involved in regulating local translation in neuronal cells^24^, accumulate in HERALD bodies (Figure 4F). Electron microscopy revealed that polysomes were detected in close proximity to, but not within, PURA-positive HERALD bodies (Figure 4G, red arrows). Together, these findings suggest that HERALD bodies represent translation-repressed, transportable mRNP granules specifically scaffolded by HSATIII RNAs, and may serve as sources of HSATIII RNAs for local translation at nearby polysomes.

**Figure 4.**
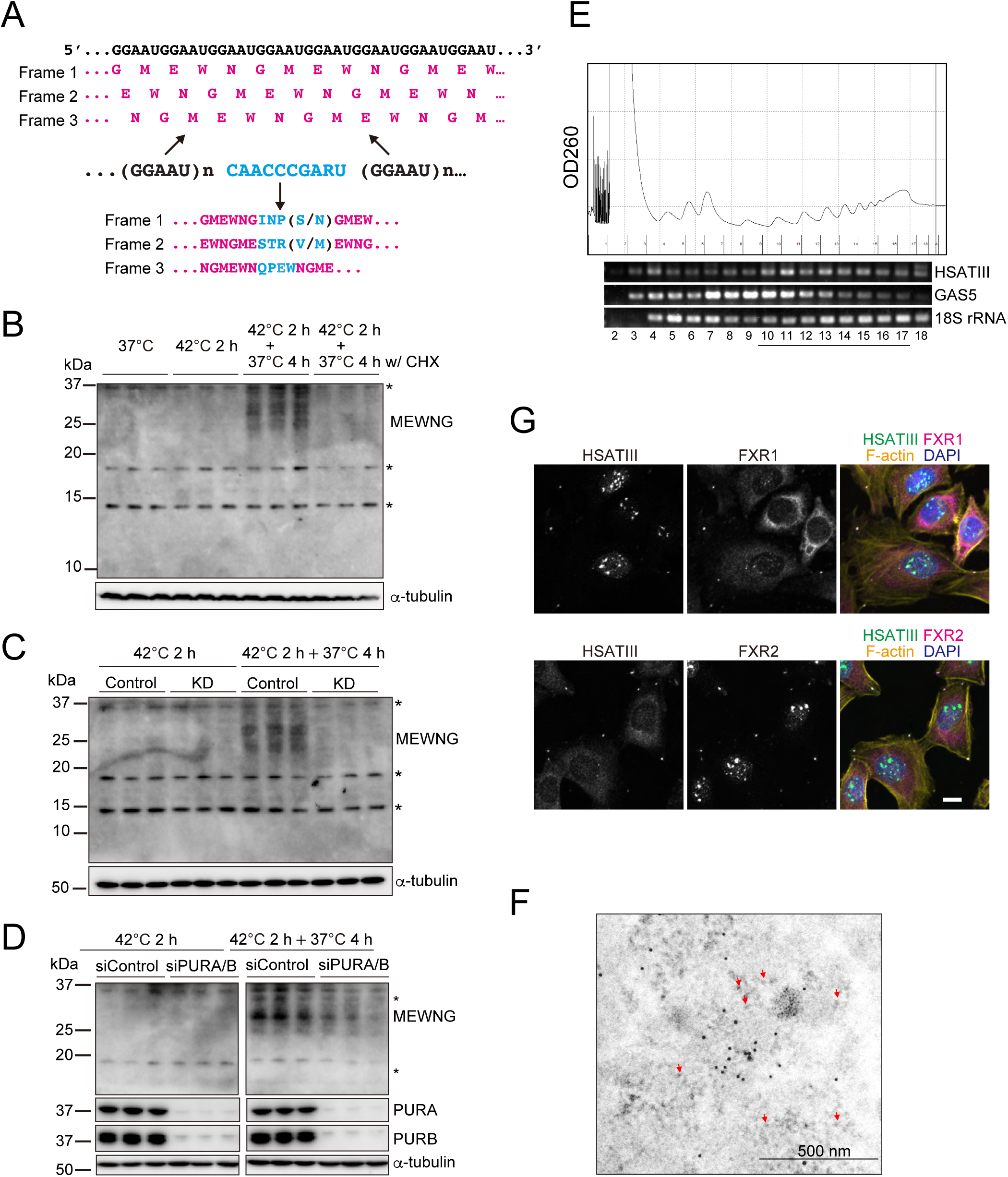
Detection of HSATIII RNA-derived proteins. (A) Theoretical amino acids sequence patterns derived from HSATIII consensus sequence. GGAAU repeat-derived polypeptide in each frame are shown in magenta. Less frequent repeat and the derived polypeptide are shown in cyan. (B) Western blot validation of HSATIII RNA-derived proteins. Control and thermal stress-exposed HeLa cells were analyzed by western blotting using anti-MEWNG antibody. a-tubulin was used as a loading control. Cycloheximide (100 mg/ml) was added during the recovery period (+CHX). Asterisks indicate non-specific bands (the same hereinafter). The data was shown as biological triplicate (The same hereinafter). (C) Validation by HSATIII KD. HeLa cells were transfected with control sense oligonucleotide (control) or HSATIII ASO (KD), cultured for 16 h and exposed to thermal stress (42℃ 2h with or without 4-hour recovery at 37℃). Western blot was performed as described above. (D) The effect of PURA/B KD on HSATIII protein production. HeLa cells were transfected with siRNAs against PURA and PURB (10 nM for each) or control siRNA (20 nM), and cultured for 48 h, and exposed to thermal stress (42℃ 2 h with or without 4-hour recovery at 37℃). Western blot was performed as described above. (E) HSATIII RNAs exist in polysome fractions. HeLa cells (42 °C for 2 h, followed by recovery at 37 °C for 2 h) were subjected to polysome fractionation analysis. An upper panel shows OD260 monitoring and lower panels show semiquantitative RT-PCR. The fraction numbers are indicated below, and the underbar indicates the polysome fractions. (F) FXR1/2 colocalization with cytoplasmic HSATIII RNAs. HeLa cells (42℃ 2 h followed by 4-hour recovery at 37℃) were stained by HSATIII-FISH and IF using anti-FXR1 or anti-FXR2 antibody. F-actin and nucleus were stained by phalloidin-iFlour^TM^647 and DAPI, respectively. Scale bar: 10 mm (G) Electron microscopy of HERALD bodies. HeLa cells (42℃ 2 h followed by 4-hour recovery at 37℃) were stained using anti-PURA antibody and observed under electron microscope. Red arrows indicate polysomes.

### ALYREF is required for the cytoplasmic functions of HSATIII RNAs

To investigate how HSATIII RNAs relocate to the cytoplasm after the stress removal, first we focused on ALYREF, a nuclear RNA export adaptor protein in TREX (TRanscription-EXport) complex^25^, because ALYREF as well as other components of the TREX complex such as DDX39A/B and CHTOP were identified as HSATIII RNA binding proteins by our HSATIII-ChIRP-MS ^11^. Notably, ALYREF was more enriched in the HSATIII-ChIRP fraction from HeLa cells 1 hour after stress removal than under thermal stress conditions (Figure 1C)^10^. Then, we hypothesized that ALYREF was transiently recruited to nSBs in the early phase of recovery to promote nuclear export of HSATIII RNAs, resulting in HERALD formation and HSATIII polypeptide synthesis in the later phase. To test the hypothesis, we performed ALYREF KD and tested the effects on cytoplasmic functions of HSATIII RNAs. In the ALYREF KD cells, both the number and size of HERALD foci were decreased (Figure 5A, B, Figure S4A) and the production of HSATIII polypeptides was also inhibited (Figure 5C). Next, we tested whether the enhanced recruitment of ALYREF into nSBs up-regulates the formation of HERALDs and the production of HSATIII proteins. We coincidentally found that KD of SAFB, one of major nSB proteins, enhanced the recruitment of ALYREF to nSBs (Figure S5A and S5B). Then, we tested whether the upregulation of ALYREF recruitment to nSBs by SAFB KD enhances the cytoplasmic functions of HSATIII RNAs. In this situation, the number of HERALD foci and the production of HSATIII proteins remarkably increased (Figure 5D and 5E). These effects by SAFB KD were largely prevented by additional KD of ALYREF (Figure 5F-H), suggesting that the effects of SAFB depletion are mediated by enhanced localization of ALYREF to nSBs. Collectively, these data suggest that both HERALD formation and HSATIII protein production during the 4-hour recovery phase depend on ALYREF, which is recruited to nSBs at the onset of recovery.

**Figure 5.**
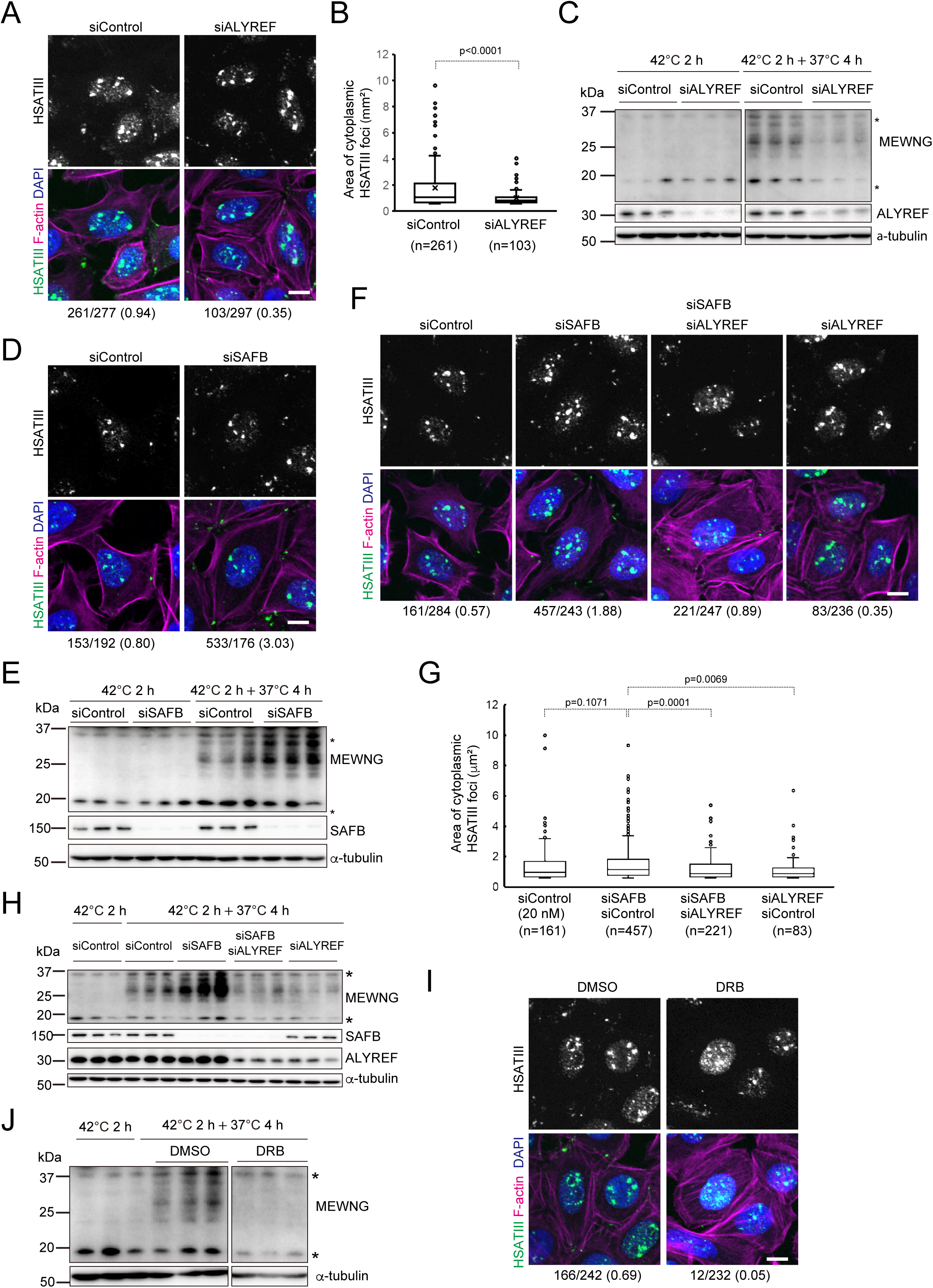
ALYREF as a trigger for cytoplasmic function of HSATIII RNAs. (A) The effect of ALYREF KD on cytoplasmic HSATIII RNA foci. HeLa cells were transfected with siRNA against ALYREF (10 nM) or control siRNA (10 nM), cultured for 48 h, exposed to thermal stress (42 °C for 2 h, followed by recovery at 37 °C for 4 h), and stained by HSATIII-FISH. F-actin and nuclei were stained by phalloidin rhodamine and DAPI, respectively. The number of cytoplasmic foci per that of nuclei obtained from at least three microscopic images is indicated (the same hereinafter). Scale bar: 10 mm (B) Box plot of area of individual cytoplasmic HSATIII foci. The numbers of quantified foci are indicated below. p-value (Mann–Whitney U-test) is shown above. (C) The effect of ALYREF KD on the HSATIII protein production. HeLa cells were transfected with siRNAs against ALYREF (10 nM) or control siRNA (10 nM), and cultured for 48 h, exposed to thermal stress (42℃ 2 h with or without 4-hour recovery at 37℃) and analyzed by western blotting using anti-MEWNG antibody. a-tubulin was used as a loading control. The data was shown as triplicate. Asterisks indicate non-specific bands (the same hereinafter). (D) The effect of SAFB KD on cytoplasmic HSATIII foci formation. HeLa cells were transfected with siRNA against SAFB or control siRNA (10 nM), and cultured and stained as described in (A). Scale bar: 10 mm (E) The effect of SAFB KD on HSATIII protein production. HeLa cells were transfected with siRNA against SAFB or control siRNA (10 nM), and cultured and analyzed as described in (C) (F) The effect of combinational KD of SAFB and ALYREF. HeLa cells were transfected with control siRNA (20 nM), SAFB and control siRNAs (10 nM each), SAFB and ALYREF siRNAs (10 nM each) or ALYREF and control siRNAs (10 nM each), cultured and stained as described in (A). (G) Box plot of area of individual cytoplasmic HSATIII foci (0.5 - 10 mm^2^) in (F). The numbers of quantified foci are indicated below. P-Value (Kruskal-Wallis test, followed by Dunn’s multiple comparison test) is shown above (H) The effect of combinational KD of SAFB and ALYREF on the HSATIII protein production. HeLa cells were prepared as described in (F) and analyzed by western blotting as described in (C) (I, J) The effects of DRB treatment during the recovery on the HERALD formation (I) and HSATIII protein production (J). HeLa cells were exposed to thermal stress (42 °C for 2 h), cultured at 37 °C for 4 h in the presence of DMSO or DRB (100 mM), and analyzed as described in (A) and (C).

Transcription of HSATIII RNAs is induced by HSF1 activation upon thermal stress^26^ and continues for at least within 1 hour after stress removal^11^. Consistently, HSF1 remains accumulated in nSBs for at least 1 hour after stress removal but becomes diffuse by 4 hours (Figure S5C). These observations collectively suggest that HSATIII transcription persists for several hours into the stress period. Then, we tested whether the cytoplasmic HSATIII RNAs are derived from pre-existing transcripts produced during the stress phase or newly synthesized RNAs during recovery. When the transcription was inhibited by DRB during the recovery period, both HERALD formation and HSATIII protein production were significantly suppressed (Figure 5I and 5J). Furthermore, ChIRP-western blotting revealed that ALYREF interaction with HSATIII RNAs was abolished in the presence of DRB, whereas the recovery phase specific interaction of TDP-43 remained unaffected (Figure S5D). These results suggest that ALYREF is recruited to nSBs through interactions with de novo transcribed regions of HSATIII RNAs produced during the recovery phase, thereby mediating their nuclear export.

### HSATIII proteins interact with HSPA8, ATXN2L and actin

To investigate the functionality of HSATIII polypeptides, we searched for their interacting proteins. We performed a comprehensive interactome analysis using immunoprecipitation coupled with mass spectrometry (IP-MS) and proximity labelling with TurboID followed by MS (TurboID-MS) as schematically shown in Figures 6A and 6B. First, we performed co-IP using anti-MEWNG antibody and identified coprecipitated proteins by MS. To improve IP efficiency, we utilized SAFB KD cells, in which HSATIII protein production was enhanced (Figure 5E). In addition, to minimize nonspecific binding, we conducted parallel IP-MS experiments using SAFB KD cells under HSATIII-induced conditions (42℃ for 2 h followed by recovery at 37℃ for 4 h; induced in Figure 6A) and non-induced conditions (37℃; control in Figure 6A). Proteins more efficiently precipitated from the induced sample were selected (enrichment score ratio (induced /control) >3) (Figure 6A, Table S1). As additional negative control, we also performed IP-MS using normal rabbit IgG, and selected proteins specifically precipitated by the anti-MEWNG antibody (enrichment score ratio (anti-MEWNG /normal IgG) >3) (Figure 6A, Table S1). At the same time, we performed Turbo-ID-MS screening. Since HSATIII proteins are likely heterogeneous, originating from various satellite III genomic regions, and the amino acid sequences of each protein have not yet been determined, we constructed TurboID-fused artificially-designed HSATIII proteins (adHSATIII proteins shown in Figure S6A). To construct artificial HSATIII sequences, we performed PCR-based repeat extension of predominantly GGAAU repeat sequences with or without CAACCCGAAU insertions at regular intervals. We obtained various patterns of long DNA fragments containing artificial HSATIII-like repeat sequences, although they contained a few mutations and sequence-undetermined regions due to technical limitations for the fidelity of repeat extension PCR and the difficulty in sequencing long repetitive DNA (Figure S6A and Table S2). Then, we constructed two TurboID expression vectors containing artificial HSATIII proteins (adHSATIII); one expressing a MEWNG-based repeat polypeptide (∼33-kDa; adHSATIII#8), and the other expresses a 33-kDa polypeptide in which MEWNG repeats are interspersed with CAACCCGAAT-derived amino acids (adHSATIII#32). TurboID-MS was performed using HeLa cells expressing TurboID-fused adHSATIII proteins or TurboID alone as a negative control, and proteins labeled in proximately to adHSATIII proteins were identified (enrichment score ratio (TurboID-adHSATIII protein #8 or #32 /TurboID alone) >2) (Table S1). We obtained 31 proteins that were positive in all four screening assays (Figure 6C). To minimize the risk of false negatives, we additionally included 155 proteins that tested positive in three of four assays. Notably, 186 candidate proteins included stress granule components such as ATXN2L and ZC3HAV1(also referred to as ZAP) and heat shock proteins such as HSPA8 (also referred to as HSC70). Western blot validation of the IP samples confirmed that these proteins were co-precipitated with endogenous HSATIII proteins (Figure 6D and 6E). They were also co-precipitated with various types of FLAG-tagged HSATIII proteins derived from an endogenous HSATIII fragment (clone #10; Figure 6F and Figure S6B) and artificial HSATIII proteins (Figure S6A and S6C). HSPB1 (HSP27), another candidate among HSP family proteins, also interacted with FLAG-tagged HSATIII proteins (Figure 6F and 6SC). These results indicate that these proteins commonly interact with a broad range of HSATIII proteins despite variations in their lengths and amino acid sequences. We also identified ACTB (beta-actin) and F-actin capping protein, CAPZ as HSATIII-interacting proteins (Figure 6D-F, S6C), which will be described later in more detail.

**Figure 6.**
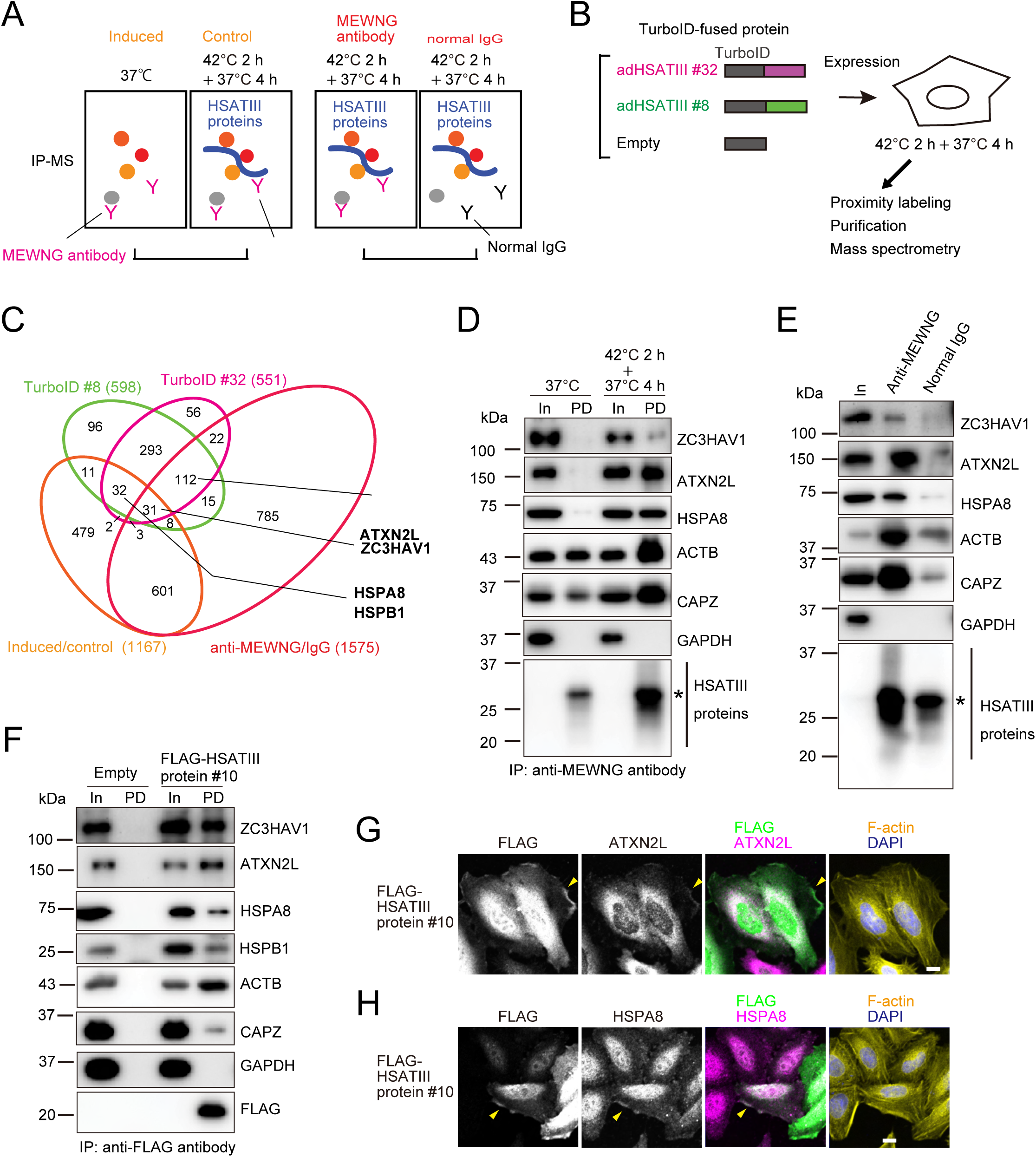
Interactome of HSATIII proteins. (A, B) Schemes of IP-MS (A) and TurboID-MS (B) screenings for HSATIII protein binding proteins. (C) Summary of HSATIII protein interactome. Venn’s diagram shows overlapping of proteins identified by IP-MS using anti-MEWNG antibody (red and orange) and TurboID using artificial HSATIII proteins #8 (green) and #32 (magenta). Proteins analyzed in the following assays are indicated. (D-F) IP-western blot validation of HSATIII protein binding proteins. IP was performed using anti-MEWNG antibody from HeLa cells (37 °C or 42 °C 2 h + 37 °C 4 h) (D), using anti-MEWNG antibody or normal rabbit IgG from HeLa cells (42 °C 2 h + 37 °C 4 h) (E), and using anti-FLAG antibody from HeLa cells (42 °C 2 h + 37 °C 4 h) expressing exogenous FLAG-tag HSATIII protein #10 (F). Western blot analysis was performed using indicated antibodies. GAPDH was used as negative control. In (Input 1%), PD (pulldown). Asterisks show signals derived from contaminated IgG light chain and/or protein G. (G, H) Colocalization of HSATIII proteins and the binding proteins at the cell periphery. Hela cells expressing FLAG-tag exogenous HSATIII protein #10 were exposed to thermal stress (42 °C 2 h + 37 °C 4 h) and stained by IF using antibodies against FLAG-tag and ATXN2L (G) or HSPA8 (H). F-actin and nucleus were stained by phalloidin-iFlour^TM^647 and DAPI, respectively. Scale bar: 10 mm

### HSATIII proteins form actin cytoskeleton-associated protein complexes with HSPA8 and ATXN2L

Next, we investigated whether the interacting proteins identified above colocalize with HSATIII proteins. First, we examined the subcellular localizations of ATXN2L and HSPA8 before and after thermal stress. HSPA8 was diffusely distributed in the cytoplasm under normal conditions and was localized in the nucleus during thermal stress (Figure S6D) as previously reported ^27,28^. ATXN2L also diffused in the cytoplasm under normal conditions but exhibited SG-like assemblies during thermal stress (Figure S6D), consistent with previous findings^29^. Interestingly, we found that 4 hours after stress removal, during the recovery phase, these proteins partially colocalized with the actin cytoskeleton at the cell periphery (Figure S6D, arrowheads). To examine whether HSATIII proteins colocalize with these factors, we used exogenously expressed FLAG-tagged HSATIII proteins because the anti-MEWNG antibody used for western blotting of HSATIII proteins is unfortunately not suitable for immunostaining of endogenous HSATIII proteins due to strong nonspecific signals. FLAG-tagged and artificially designed HSATIII proteins were broadly distributed in both the nucleus and the cytoplasm, but colocalized with ATXN2L and HSPA8 at the cell periphery (Figure 6G, 6H, S6D and S6F, arrowheads), consistent with the interaction of HSATIII proteins with actin cytoskeleton proteins (ACTB and CAPZ), as well as HSPA8 and ATXN2L (Figure 6D-F and S6B). We also found that actin polymerization induced by the actin filament-stabilization agent, jasplakinolide caused aggregation of these proteins near the actin polymerization sites (Figure 7A and 7B, Figure S7A). ZC3HAV1, and another stress granule protein, G3BP2, and HSPB1 also accumulated at jasplakinolide-induced ATXN2L/HSPA8 foci (Figure S7B-E), indicating that these aggregates contain multiple SG proteins and HSPs. In addition, ACTB or CAPZ were not coprecipitated with FLAG-tagged HSATIII proteins in cells pre-treated with the actin-depolymerization agent latrunculin A, whereas the interactions of HSATIII proteins with ZC3HAV1, ATXN2L, HSPA8 and HSP1B were largely unaffected (Figure 7C), suggesting these interactions are independent of polymerized actin. Collectively, these results suggest that HSATIII proteins form novel complexes containing HSPs and SG factors, and that their intracellular distribution is regulated in association with actin cytoskeleton dynamics.

**Figure 7.**
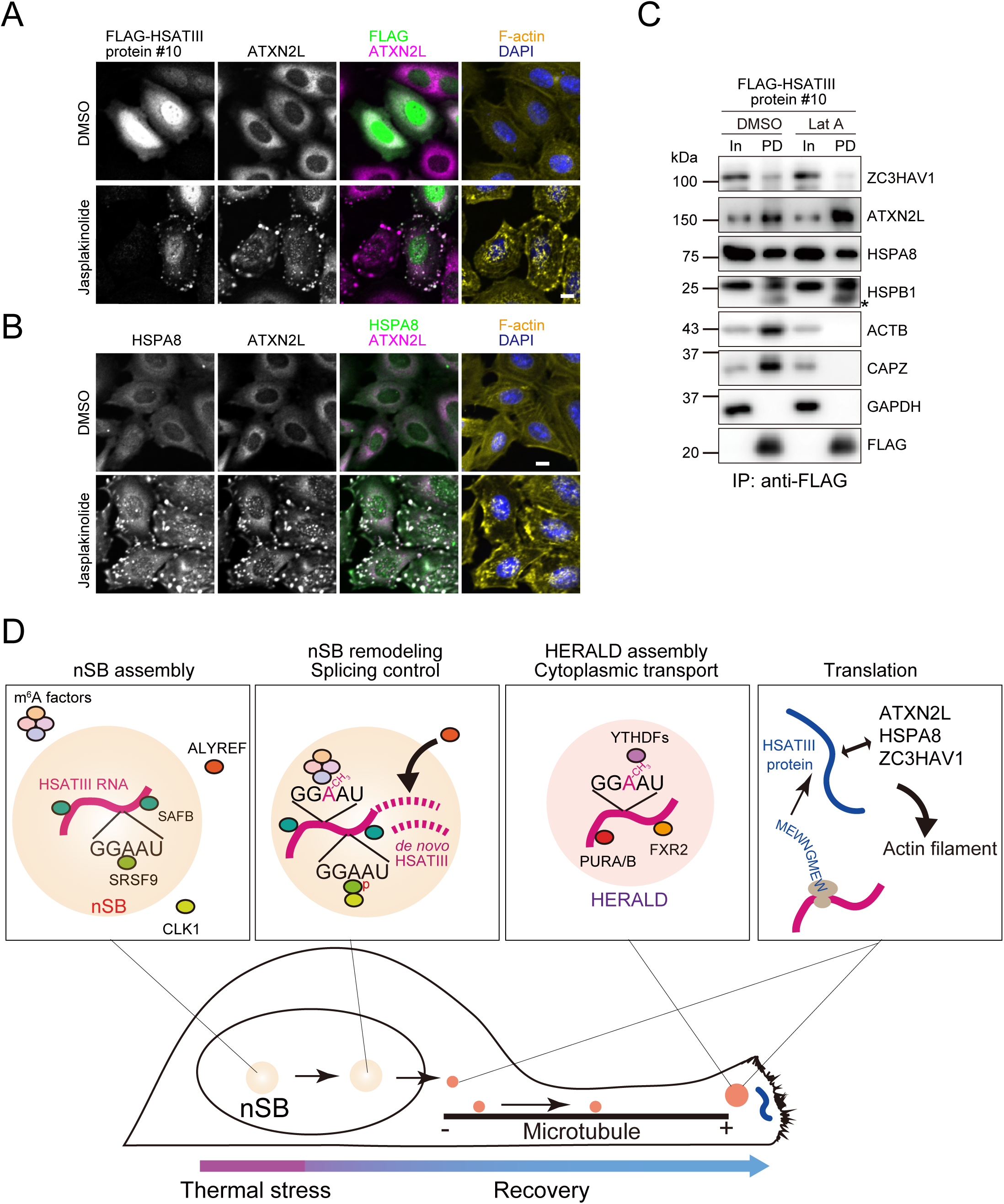
Actin polymerization-dependent aggregation of HSATIII protein complexes. (A, B) Actin polymerization-dependent aggregation of HSATIII protein complexes. HeLa cells prepared same as Figure 6G were treated with jasplakinolide (100 nM, 30 min) or DMSO as control before fixation and stained using specific antibodies. F-actin and nucleus were stained by phalloidin-iFlour^TM^647 and DAPI, respectively. Scale bar: 10 mm (D) The effect of actin-depolymerization on HSATIII protein complexes. HeLa cells expressing FLAG-tagged HSATIII protein #10 were exposed to thermal stress (42 °C for 2 h, followed by recovery at 37 °C for 4 h). The cells were treated with DMSO or latrunculin A (Lat A, 200 ng/ml, 30 min), lysed in Pierce lysis IP buffer containing DMSO or Lat A and analyzed by IP-WB as described in Figure 6F. (E) Model for structural and functional remodeling of HSATIII-based RNP bodies.

## DISCUSSION

HSATIII RNAs have been recognized to function as architectural lncRNAs that regulate splicing by driving the assembly of nSBs in the nucleus^9^. In this study, we demonstrate additional molecular roles of HSATIII RNAs in the cytoplasm (Figure 7D). Specifically, during the recovery phase after thermal stress, HSATIII RNAs relocate to the cytoplasm and form novel RNP granules together with PURA, PURB, TDP-43, FXR1, FXR2 and cytoplasmic m^6^A reader proteins, YTHDFs. These RNP granules, termed HERALDs, accumulate primarily at the distal cellular sites, and their formation is abolished when microtubule polymerization is inhibited. Given that PURA/B interact with kinesin motor proteins^21^ and are required for HERALD formation, HERALD assembly at cell peripheries is likely driven by kinesin-mediated microtubule transport. Moreover, several HERALD protein components, including FXR1/2 and YTHDFs, are also found in other transportable RNPs that carry heterogenous mRNAs for local translation, especially in neuron^30^. Based on these similarities, we propose that HERALDs represent a specialized class of transportable mRNPs scaffolded by HSATIII RNAs, which accumulate at distal cellular sites in anticipation of stimulus-induced local translation. Although we have not yet detected distal-targeted mRNAs such as APC in HSATIII-ChIRP fractions, we cannot exclude the presence of other mRNAs within HERALDs. An intriguing possibility is that cytoplasmic HSATIII RNAs facilitate HERALD formation to promote distal targeting of specific mRNAs, or alternatively, impede cytoplasmic trafficking of such mRNAs by sequestrating key transport factors.

Our data demonstrate that recovery-phase-specific recruitment of ALYREF on HSATIII RNAs triggers their nuclear export, leading to HERALD formation and HSATIII protein production, although it remains unclear how ALYREF preferentially interacts with HSATIII RNAs after the temperature returns to normal. First, the ALYREF interaction and the subsequent cytoplasmic functions of HSATIII RNAs, requires de novo transcription during the recovery period. This raises an intriguing possibility that HSATIII RNAs form distinct RNPs and follow different fates depending on when they are transcribed, during thermal stress or after the stress removal. TDP-43 also interacts with HSATIII RNAs specifically during the recovery phase. However, unlike ALYREF, its interaction is not affected by transcription inhibition (Figure S5D), suggesting that the recovery-depending engagement of TDP-43 occurs post-transcriptionally.

Second, the interaction of ALYREF with HSATIII RNAs and its localization within nSBs were enhanced upon SAFB KD, suggesting that SAFB competitively inhibits ALYREF association with HSATIII RNAs. This raises the possibility that the relative affinities of SAFB and/or ALYREF for newly transcribed HSATIII RNAs may shift after thermal stress is removed. nSBs are known to form during thermal stress and undergo remodeling during recovery. For example, CLK1 kinase is excluded from nSBs during stress but recruited after the stress removal, where it phosphorylates SRSFs and promotes intron detention of various mRNAs^11^. In parallel, HSATIII RNAs become m^6^A-modified during recovery and recruit YTHDC1 within nSBs, thereby competitively inhibiting m^6^A-dependent splicing of other pre-mRNAs^10^. Notably, SRSFs and YTHDC1 have been reported to bind ALYREF and promote its interaction with cargo mRNAs^31^. Therefore, SRSF phosphorylation and/or m^6^A modification events occurring within nSBs after stress removal may drive ALYREF recruitment to HSATIII RNAs. Further studies will be required to determine whether, and how, these remodeling-associated modifications cooperatively regulate the localization and functions of HSATIII RNAs.

We also show that HSATIII RNAs are translated into heterogenous HSATIII proteins. Determining the precise amino acid sequences of these proteins is challenging due to the heterogeneity in length and sequence of HSATIII repeat RNAs and the technical difficulties in sequencing highly repetitive RNAs. Furthermore, the open reading frames (ORFs) and utilized initiation codons are difficult to predict. The GGAAU repeat motif of HSATIII RNAs contains numerous AUG triplets, making it unclear whether translation initiates at the first AUG or at downstream AUGs. Non-AUG RAN translation also cannot be ruled out as an additional translational mechanism. Regardless of the initiation mode, most translation products are expected to share a common MEWNG motif encoded by the GGAAU repeat (Figure 4A). In this study, we used endogenous and/or artificial HSATIII proteins to analyze their interactome by IP-MS and TurboID-MS and to assess their subcellular localization. These analyses revealed that HSATIII proteins form complexes with ATXN2L, ZC3HAV1, HSPA8, and HSPB1 at the cell periphery, where they closely associate with the actin cytoskeleton specifically during the recovery phase after thermal stress. Notably, diverse HSATIII polypeptides examined here interacted and colocalized with these proteins and with actin, suggesting that the interactions are robust to variations in polypeptide sequence and length. HSPA8 has been shown to bind the actin-capping protein CAPZ^33^, and HSP1B has been proposed to associate with F-actin and inhibit actin polymerization in vitro^34–36^. However, to our knowledge, the formation of an actin-associated complex containing HSATIII proteins and ATXN2L during stress recovery is novel. Given that ZC3HAV1, HSPA8 and ATXN2L have been implicated in invasion or degradation of viral RNAs^37–40^, these HSATIII protein complexes may influence pro- or antiviral responses. Similar repeat polypeptides are detected in Purikinje cells of SCA31 (Spinocerebellar ataxia type 31) patients^32^, SCA31 is classified into repeat expansion diseases and these polypeptides (referred to as WNGME pentapeptide repeat) are derived from pathogenic expanded GGAAU repeat in intron 6 of *BEAN1* transcript, while their molecular function and role in pathogenesis are still unclear. Our findings will also provide insights into the pathogenic mechanism of SCA31. Nevertheless, their precise biological roles remain unclear. Further investigation will be crucial to establish the functional significance of these complexes at the cell periphery, especially in the context of thermal stress and recovery.

### Limitations of the study

Because HSATIII RNA is composed of highly repetitive sequences and transcribed from multiple genomic loci, it is difficult to attribute specific functions to individual HSATIII transcripts. Moreover, since cytoplasmic HSATIII RNA cannot be completely separated from the nuclear fraction, HERALD-associated and nSB-associated HSATIII RNAs cannot be reliably distinguished by sequencing analysis.

At the protein level, the repetitive nature of the RNA and the abundance of AUG codons hinder accurate identification of the translation initiation site. Additionally, since the available HSATIII antibody is applicable only to Western blotting and immunoprecipitation but not to immunostaining, the subcellular localization of endogenous HSATIII-derived translation products cannot be determined. Consequently, the relationship between HERALD formation and local translation remains unresolved.

## RESOURCE AVAILABILITY

### Lead contact

Further information and requests for resources and reagents should be directed to and will be fulfilled by the Lead Contact, Tetsuro Hirose.

### Materials availability

All unique plasmids and constructs generated in this study are available from the Lead Contact.

### Data and code availability

All data reported in this paper will be shared by the Lead Contact upon reasonable request. The mass spectrometry raw data were deposited in the Japan ProteOme STandard Repository (jPOST) under the ID JPST. No original code was generated

## ACKNOWLEDGMENTS

We would like to thank the members of the Hirose laboratory for their valuable discussions and support. We thank FBS Core Facility at The University of Osaka for technical support. This research was supported by grants from JST CREST grant no. JPMJCR20E6 (to T.H.), JPMJCR23B3 and JPMJCR20E6 (to S.A.), AMED grant no. 21479280 (to T.H.), 23gm1610010h0002 (to S.A.), JSPS KAKENHI grant no. 21H05276 and 24K21933 (to T.H.), 22H02225 (to S.A.), 21H05277 and 25H00007 (to T.I.) National Cancer Center Research and Development Funds (2023-A-01).

## AUTHOR CONTRIBUTIONS

K.N., and T.H. conceived and designed this study. K.N. conducted most of the experiments. S.A. performed the mass spectrometry analyses. T.S. and T.I. performed polysome analysis. S.S and G.P, performed electron microscopy. K.N., and T.H. wrote the manuscript. All authors have reviewed the manuscript.

## SUPPLEMENTAL INFORMATION

### Document S1. Figures S1-S7 and Supplemental References (PDF)

**Table S1., interactome of HSATIII proteins related to Figures 6 (Excel file)**

Excel sheets show the results of IP-MS and TurboID-MS. Enrichment score ratio (A/B) of each protein was calculated as [(the average number of peptides in sample A)+0.0001]/[(the average number of peptides in sample B)+0.0001].

**Table S2., DNA sequences of endogenous and artificially designed HSATIII clones (PDF).**

**Table S3. The list of primers used in this paper, related to STAR Methods (PDF)**

The sequences of primers used for RT-PCR, cloning and repeat extension PCR in this study.

## STAR★METHODS

### KEY RESOURCES TABLE

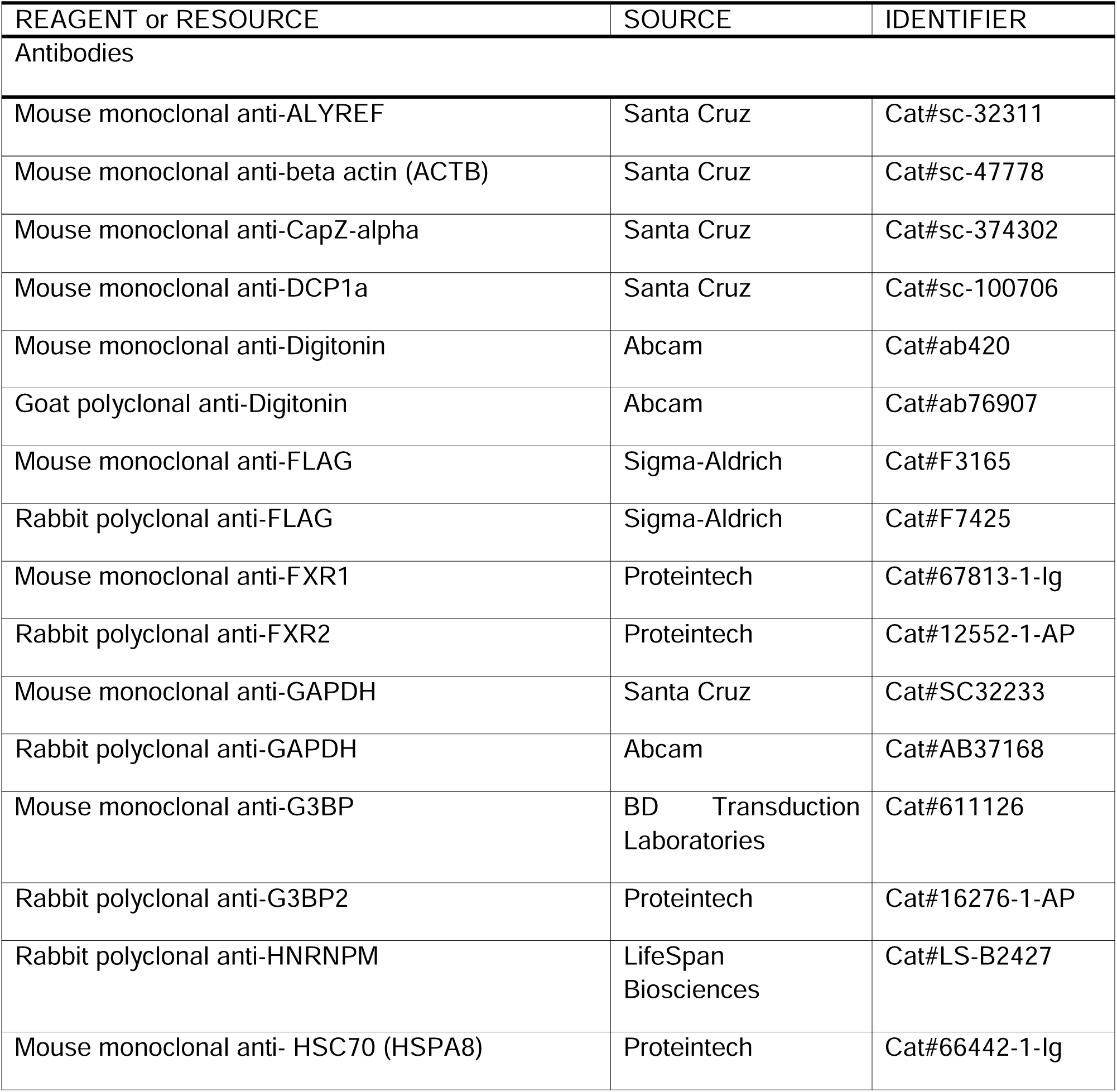
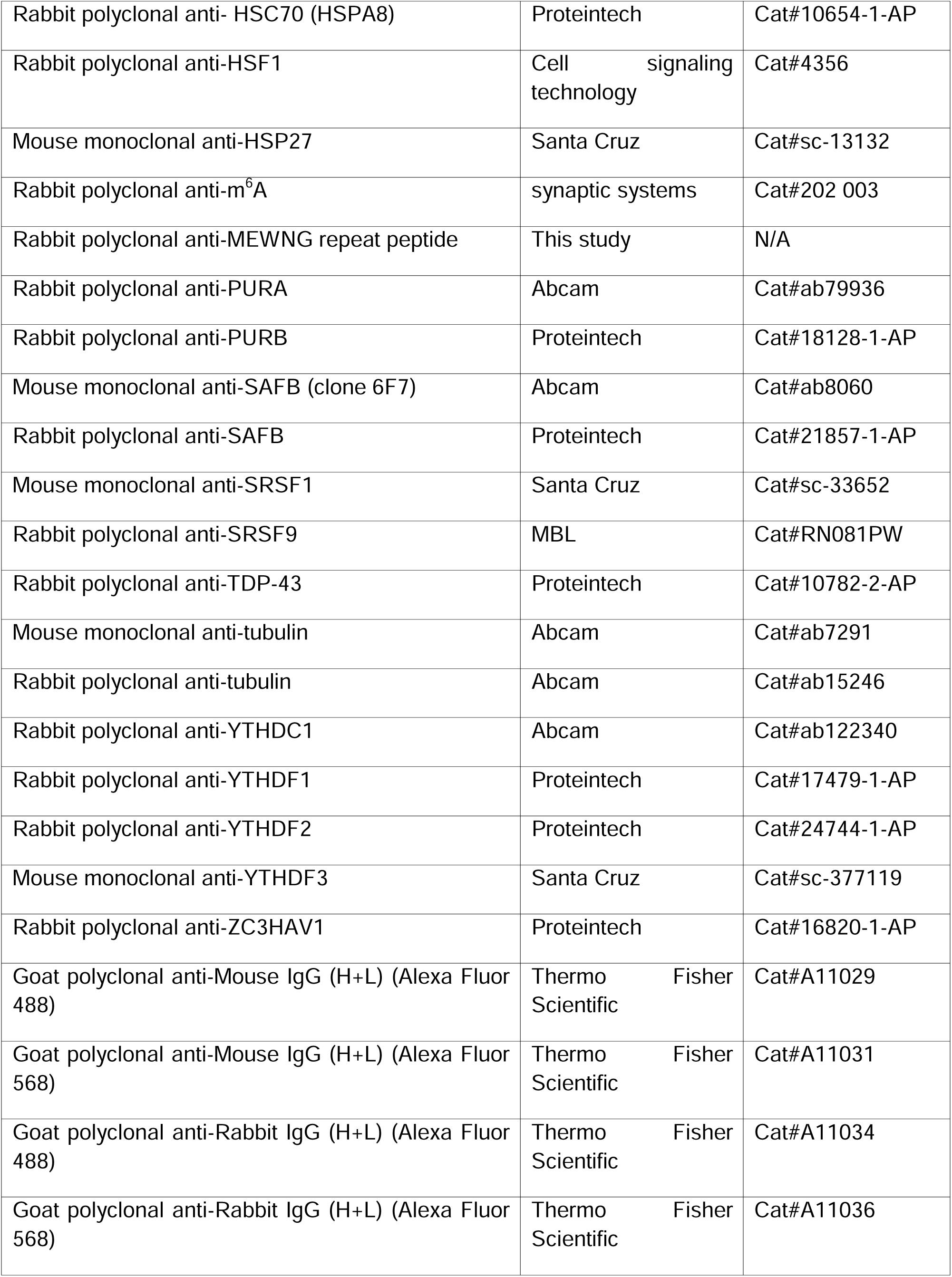
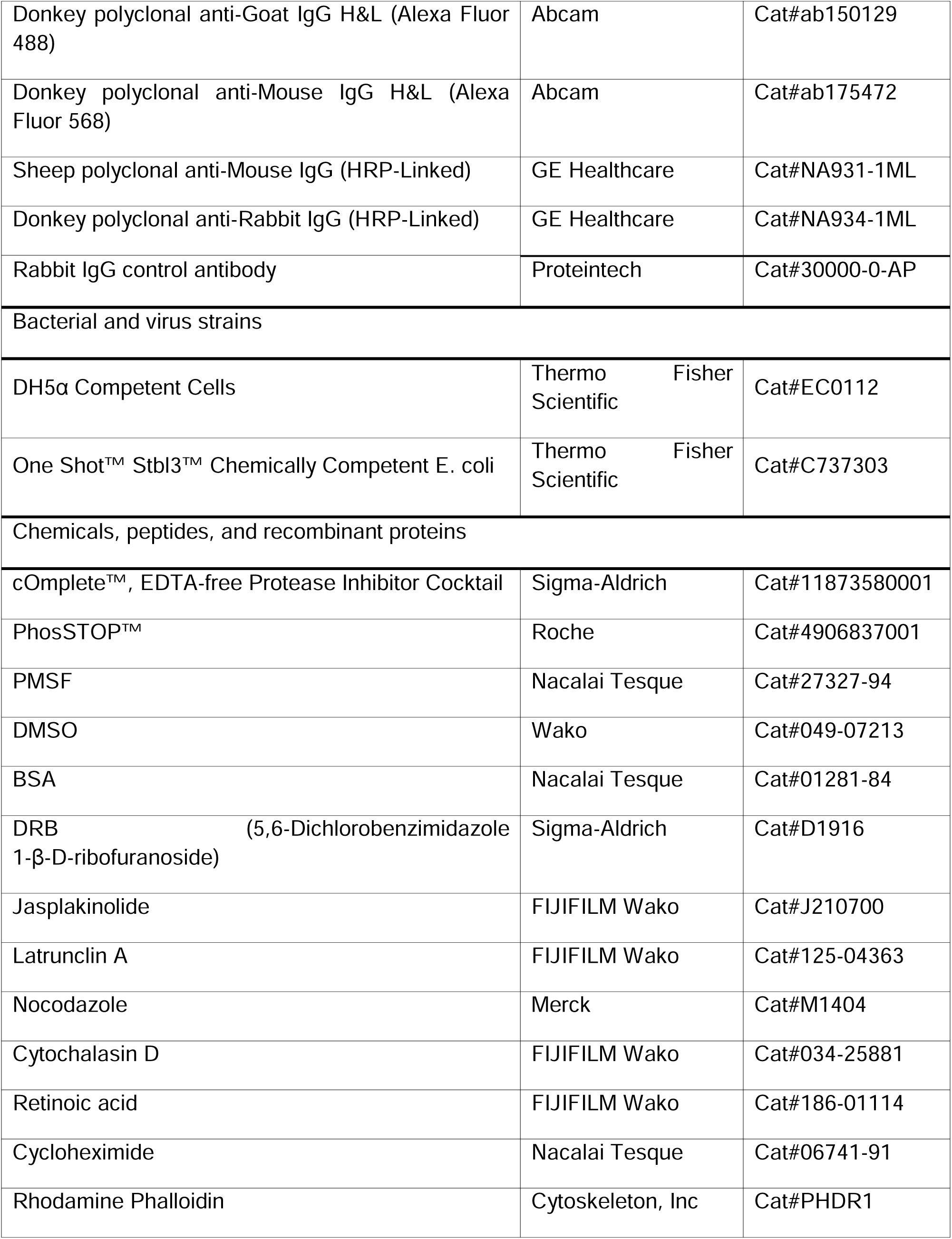
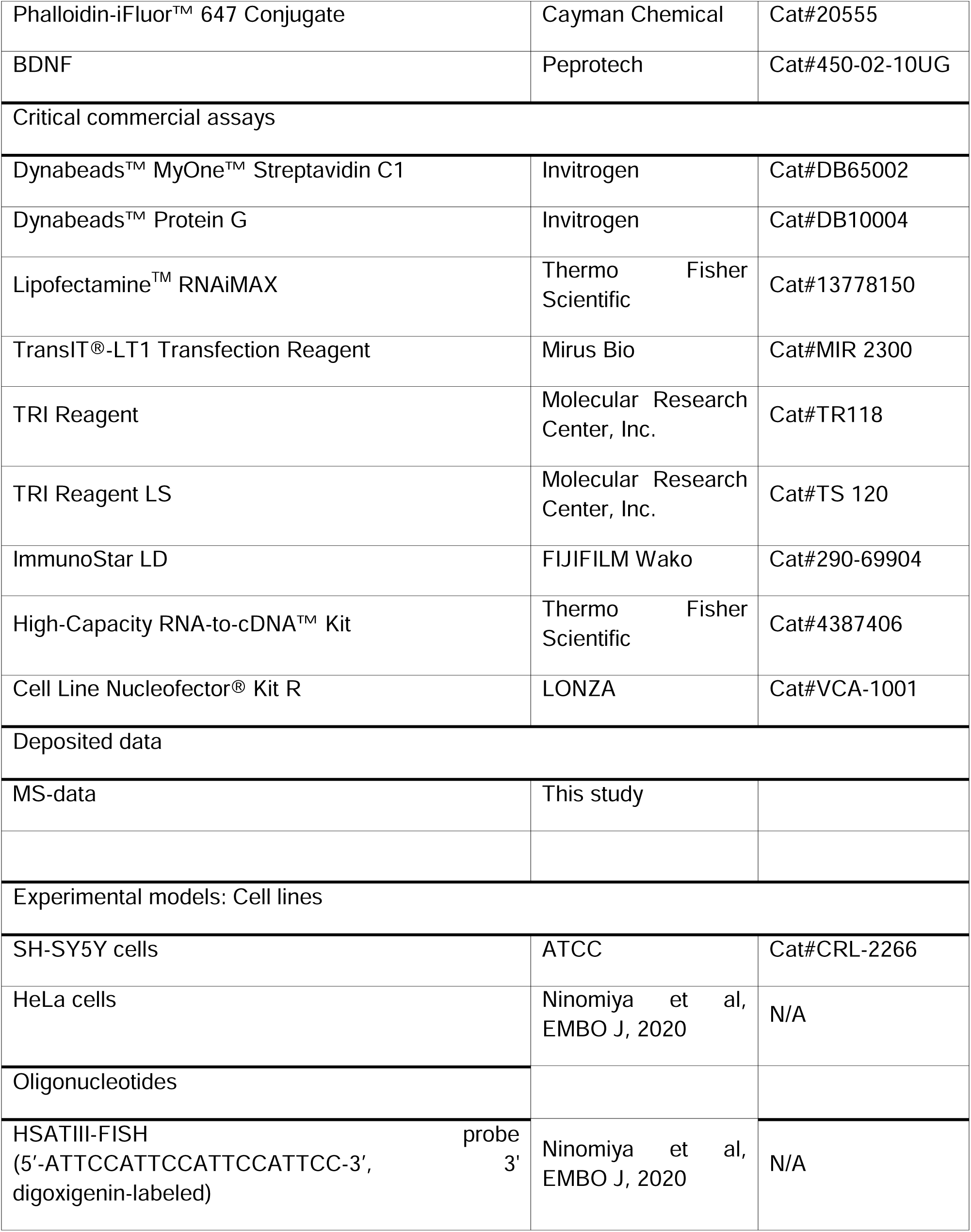
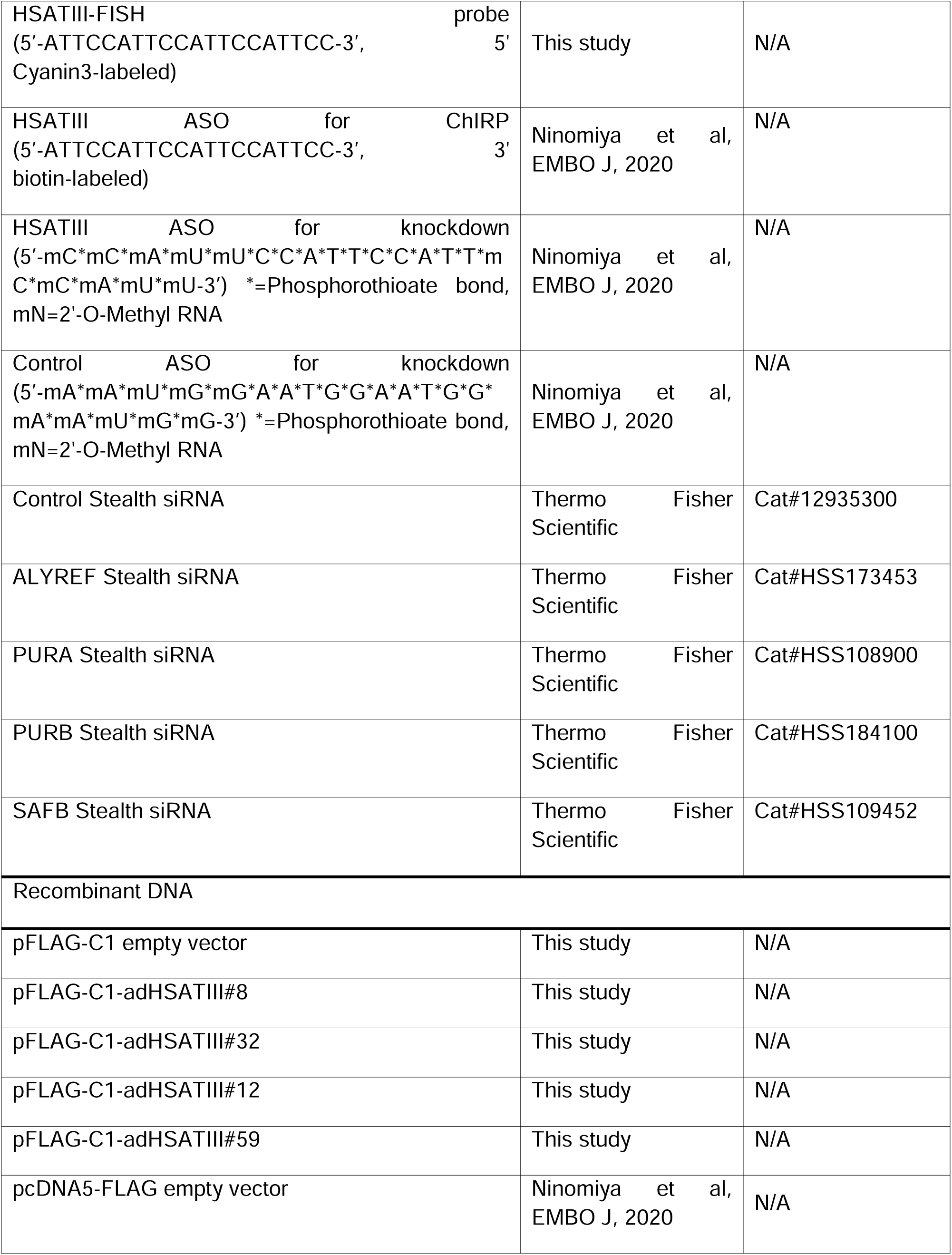
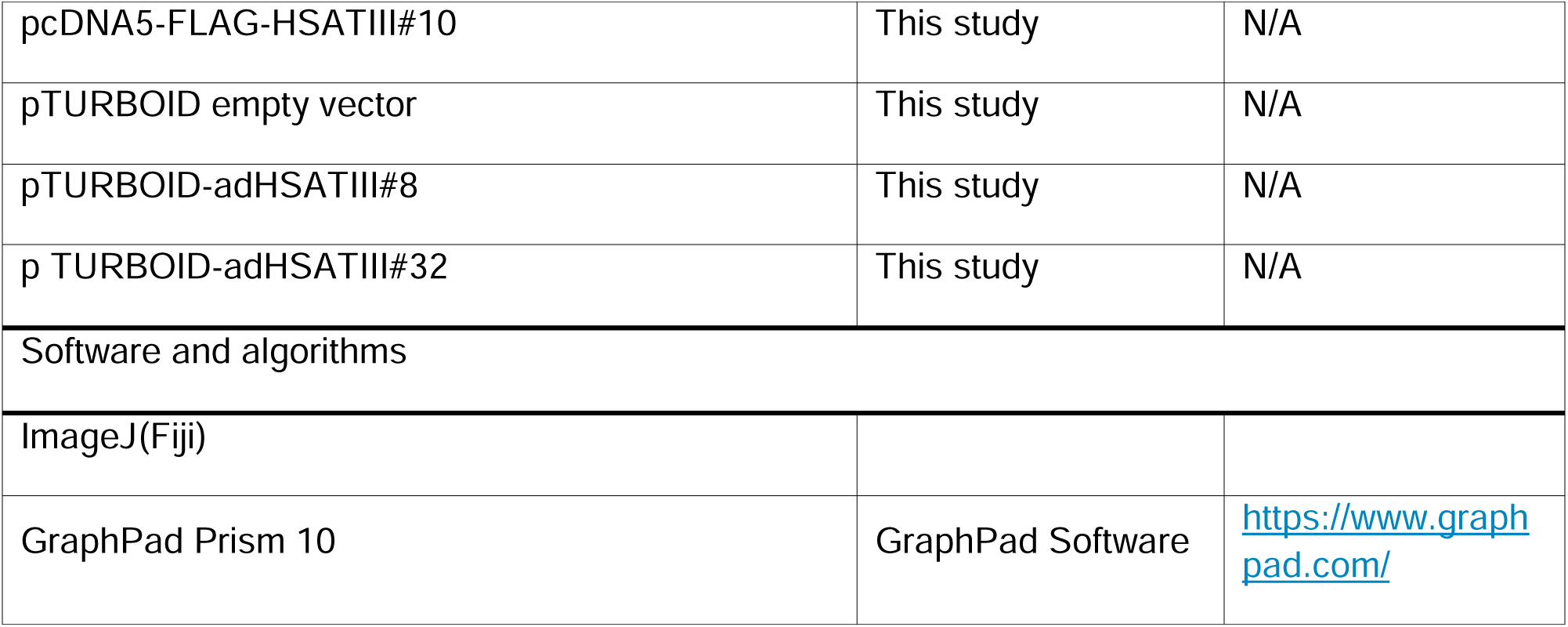

### METHOD DETAILS

#### Cell culture

HeLa cells were cultured in Dulbecco’s modified Eagle’s medium (DMEM) (Nacalai Tesque) containing antibiotics (PS; 100 U/ml streptomycin and 100 □g/ml penicillin; Sigma) and 10% fetal calf serum (Sigma) at 37°C in 5% CO_2_. For neuronal differentiation of SH-SY5Y cells, the cells (10,000 cells/cm^2^) were cultured in 10%/FBS DMEM/HAM F12 medium (Nacalai Tesque) on collagen-coated dishes (day 0) and treated with 10 □M retinoic acid (Fujifilm WAKO) from day 1 to 5 (change medium every 2 days). And then, the cells were washed three times with serum-free DMEM (Nacalai Tesque) and cultured in serum-free DMEM containing PS and 50 ng/ml recombinant human BDNF (Peprotech) from day 5 to 7-9. For thermal stress induction, culture dishes were incubated at 42°C in an incubator with 5% CO_2_.

#### Fluorescence in situ hybridization and immunofluorescence

Fluorescence in situ hybridization and immuno fluorescence were performed as previously reported^10^ with minor modifications. HSATIII RNA was hybridized with dig-labeled or Cy5-labeled HSATIII ASO. The confocal mages were obtained with FLUOVIEW FV1000 (Olympus) or NIKON FX (NIKON) and analyzed using a ImajJ fiji software. Probes and antibodies used are listed in key resource table.

#### Electron microscopy

Ultrastructural studies were carried out on ultra-thin sections of Lowicryl K4M-embedded cell pellets as previously described^41^. HeLa cells were fixed in situ with 4% formaldehyde (Electron Microscopic Sciences), scraped off from culture dishes, and centrifuged. Cell pellets were equilibrated in 30% methanol and deposited in a Leica EM AFS2/FSP automatic reagent handling apparatus (Leica Microsystems). Lowicryl polymerization under UV was performed for 40 h at -20°C and 40 h at +20°C. For immunogold electron microscopy (I-EM), ultra-thin sections were incubated at room temperature for 1 h with the primary antibody and for 30 min with the secondary anti-rabbit IgG antibody coupled to 10 nm gold particles (BBInternational). Thin sections were briefly contrasted with uranyl acetate and analyzed with a Tecnai Spirit (FEI). Digital images were taken with a SIS MegaviewIII charge-coupled device camera (Olympus).

#### ChIRP

ChIRP was performed as described previously ^10^. In brief, control and thermal stress-exposed HeLa cells were collected after crosslinking in 1% PFA/PBS for 30 min at RT. The cell pellets were lysed, sonicated and centrifuged. The supernatants were hybridized with biotinylated HSATIII antisense oligos and captured with Dynabeads MyOne Streptavidin C1 (Thermo Fisher Scientific). Bound proteins were eluted with SDS sample buffer and bound RNAs were de-crosslinked by proteinase K treatment and heating.

#### Semi-quantitative RT-PCR

Semi-quantitative RT-PCR was performed as described previously ^10^. Primers used are listed in Table S3

#### Plasmid construction

HSATIII DNA fragment was amplified by PCR, digested by restriction enzymes BamHI and NotI and introduced in frame into a pcDNA5-FLAG vector. For construction of adHSATIII expression vectors, artificially designed HSATIII-like repeat was extended by PCR-based repeat extension method using primestar GXL (TaKaRa) and cloned into pCR-Blunt II-TOPO vector (Thermo Fisher Scientific). After transformation, size-selection by colony-lysis PCR and sequence analysis, the HSATIII-like repeat DNAs were digested using BamHI and XhoI and introduced into a FLAG-tag expression vector, which was constructed by replacing the EGFP region of pEGFP-C1 vector by 3XFLAG cDNA. For construction of TurboID vectors, the TurboID DNA fragment was amplified by PCR and digested by restriction enzymes, XbaI and XhoI, and introduced in frame into adHSATIII expression vectors above digested by NheI and XhoI. PCR primers are listed in Table S3.

#### Transfection and knockdown

For siRNA knockdown, Stealth siRNA (Thermo Fisher Scientific) was transfected using Lipofectamine RNAiMAX (Thermo Fisher Scientific) 48 h before the assays. The final concentration of siRNA is described in each figure legend. For HSATIII knockdown, a modified antisense or control oligonucleotide (1 □M) was transfected into 1-3 × 10^6^ cells using Nucleofector technology (Lonza) according to the manufacture’s protocol 16–18 h before the experiments. For protein expression, an expression vector was transfected using TransIT-LT1 transfection reagent (Mirus) 16-24 h before the experiments. ASOs and siRNAs are listed in Key Resource Table.

#### Western blotting

Western blot was performed as previously reported^10^. In brief, the samples in 1× SDS sample buffer were boiled for 5 min, separated by SDS-PAGE, and transferred to a PVDF membrane (Millipore) by electroblotting. The membrane was incubated with primary and HRP-conjugated secondary antibodies. The signals were developed by a chemiluminescence reaction using the ImmunoStar LD (Fujifilm WAKO), detected using a ChemiDoc Touch imaging system (BioRad), and analyzed using ImageJ software (NIH). Antibodies used in this study are listed in Key Resource Table. An antibody against MEWNG polypeptide was generated by immunizing a rabbit using CMEWNGMEWNGMEWNGMEWNG polypeptide as antigen and purified using antigen-conjugated affinity column (Cosmo Bio).

#### Immunoprecipitation

HeLa cells were washed in ice-cold PBS and suspended in ice-cold Pierce IP Lysis Buffer (25 mM Tris•HCl pH 7.4, 150 mM NaCl, 1% IGEPAL CA-630, 1 mM EDTA, 5% glycerol) containing protease inhibitor cocktail and 1 mM PMSF. The cell homogenates were incubated for 20 min at 4°C, sheared 10 times using 27G needle and centrifuged at 14000 rpm for 10 min at 4°C. The supernatant was mixed with antibody-bound protein G dynabeads (Thermo fisher scientific) and rotated at 4°C overnight. The beads were washed five times in ice-cold TBST (50 mM Tris-HCl pH 7.6, 150 mM NaCl, 0.1% Tween20) and the bound proteins were eluted in SDS sample buffer by boiling for 5 min or gently shaking at RT for 30 min.

#### Polysome preparation

HeLa cells were treated in medium containing 100 mg/mL cycloheximide for 15 min. After removing medium, the cells were collected by scraping, transferred to 1.5 mL tube, and centrifugated (3,500 rpm, 1 min, 4℃). The collected cells were suspended in 500 ml lysis buffer [20 mM Tris-HCl pH7.5, 100 mM KCl, 10 mM MgCl_2_, 0.5%v/v NP-40, 1 mM DTT, 100 mg/mL Cycloheximide, 1 mM PMSF, cOmplete™ EDTA-free Protease Inhibitor Cocktail (Roche), 40 U/mL RNase inhibitor (Promega)] and lysed by pipetting on ice. The samples were centrifuged (2,000 xg, 5 min, 4℃) and the supernatants were transferred to new 1.5 mL tubes. Sucrose density gradients (10–50% sucrose in 20 mM Tris-HCl pH 7.5, 100 mM KCl, 10 mM MgCl_2_) were prepared using a Gradient Master 108 (BioComp) and stored at 4 °C until use. The samples were added onto the sucrose gradient and separated by centrifugation with P40ST rotor (himac) at 40,000 rpm, 4°C for 90 min. After centrifugation, the polysome profiles were generated by continuous absorbance measurement at 260 nm and fractions were collected using Piston Gradient fractionator and TRIAX flow cell monitor FC-1-260 (BioComp) connected to Micro Collector AC-5700 (ATTO). RNAs were extracted using TRI Reagent LS (Molecular Research Center, Inc.) according to the manufacturer’s protocol.

#### TurboID proximity labeling

Proximity labeling using the TurboID system was performed as previously described^42^. HeLa cells were transfected with plasmids encoding TurboID-adHSATIII#8, TuroboID-adHSATIII#32 and TurboID alone, cultured for 24 h and exposed to thermal stress and recovery (42℃ 2 h and 37℃ 4 h). For proximity labeling and subsequent MS analysis, 50 μM biotin was added to the cells and the cells were incubated for 1 h. For MS analysis, the cells were lysed in a buffer containing 25 mM Tris-HCl (pH 7.5), 150 mM NaCl, 1 mM ethylenediaminetetraacetic acid (EDTA) and 1% Triton X-100. The cell lysates were homogenized by shearing using a 26G syringe needle (Nipro). After centrifugation at 10,000 g for 2 min, the supernatant was collected (cell lysate). The biotinylated proteins were captured using SoftLink Soft Release Avidin Resin (Promega). After washing the resin with a TBS buffer containing 1% Triton X-100, the biotinylated proteins were eluted by boiling at 95°C for 2 min in a buffer containing 500 mM Tris-HCl, 10% glycerol, 2% SDS, and 5% 2-mercaptoethanol.

#### Mass Spectrometry-Based Preparation, Measurement, and Data Analysis

For immunoprecipitation (IP) samples, proteins were precipitated with trichloroacetic acid (TCA), washed with chilled acetone, and dissolved in 7 M guanidine hydrochloride. The solubilized proteins were reduced with tris(2-carboxyethyl)phosphine (TCEP), alkylated with iodoacetamide (IAA) to carbamidomethylate cysteine residues, and digested sequentially with lysyl endopeptidase (Wako) and trypsin (Thermo Fisher Scientific). For phosphoproteome analysis, cells were lysed in Easy-prep™ Lysis Buffer (Thermo Fisher Scientific) supplemented with PhosSTOP™ (Roche), followed by protein precipitation with acetone. The resulting pellets were dissolved in guanidine hydrochloride and processed in the same way as the IP samples: reduced with TCEP, alkylated with IAA, and digested with lysyl endopeptidase and trypsin. Phosphopeptides were enriched using the High-Select™ Fe-NTA Phosphopeptide Enrichment Kit (Thermo Fisher Scientific) according to the manufacturer’s instructions. All resulting peptides were analyzed using an Evosep One LC system (EVOSEP) coupled to a Q-Exactive HF-X mass spectrometer (Thermo Fisher Scientific). Chromatography was performed with 0.1% formic acid in water (solution A) and 0.1% formic acid in 99.9% acetonitrile (solution B) as mobile phases. Data were acquired in data-dependent acquisition mode, selecting the top 25 precursor ions within the m/z range of 380–1500. MS/MS spectra were searched against the human Swiss-Prot protein databases using Proteome Discoverer 2.5 with the SEQUEST search engine, and filtered at a 1% false discovery rate (FDR) for peptide-spectrum matches. The data have been deposited to the jPOST repository under accession number X.

### QUANTIFICATION AND STATISTICAL ANALYSIS

Western blot, FISH and IF images were analyzed with ImageJ (FIJI). Statistical significance was assessed using GraphPad Prism 10. Statistical tests and p-values are described in figure legends and graphs.

