## Supplemental Figures S1-S7 for "Cytoplasmic roles of HSATIII RNAs in RNA granule assembly and production of actin cytoskeleton-associated repeat-containing proteins"

Supplemental figure 1

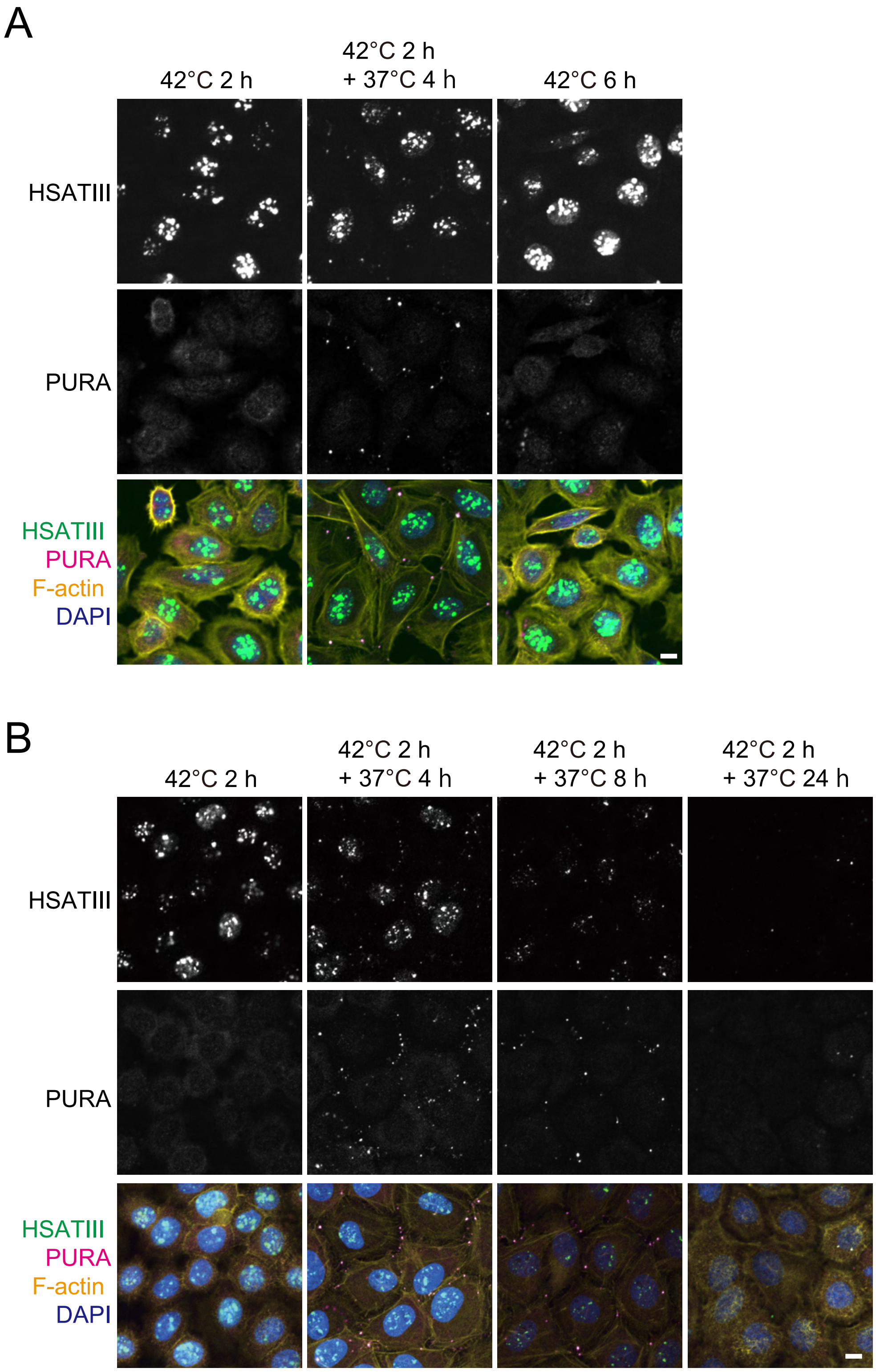

Supplemental figure 2

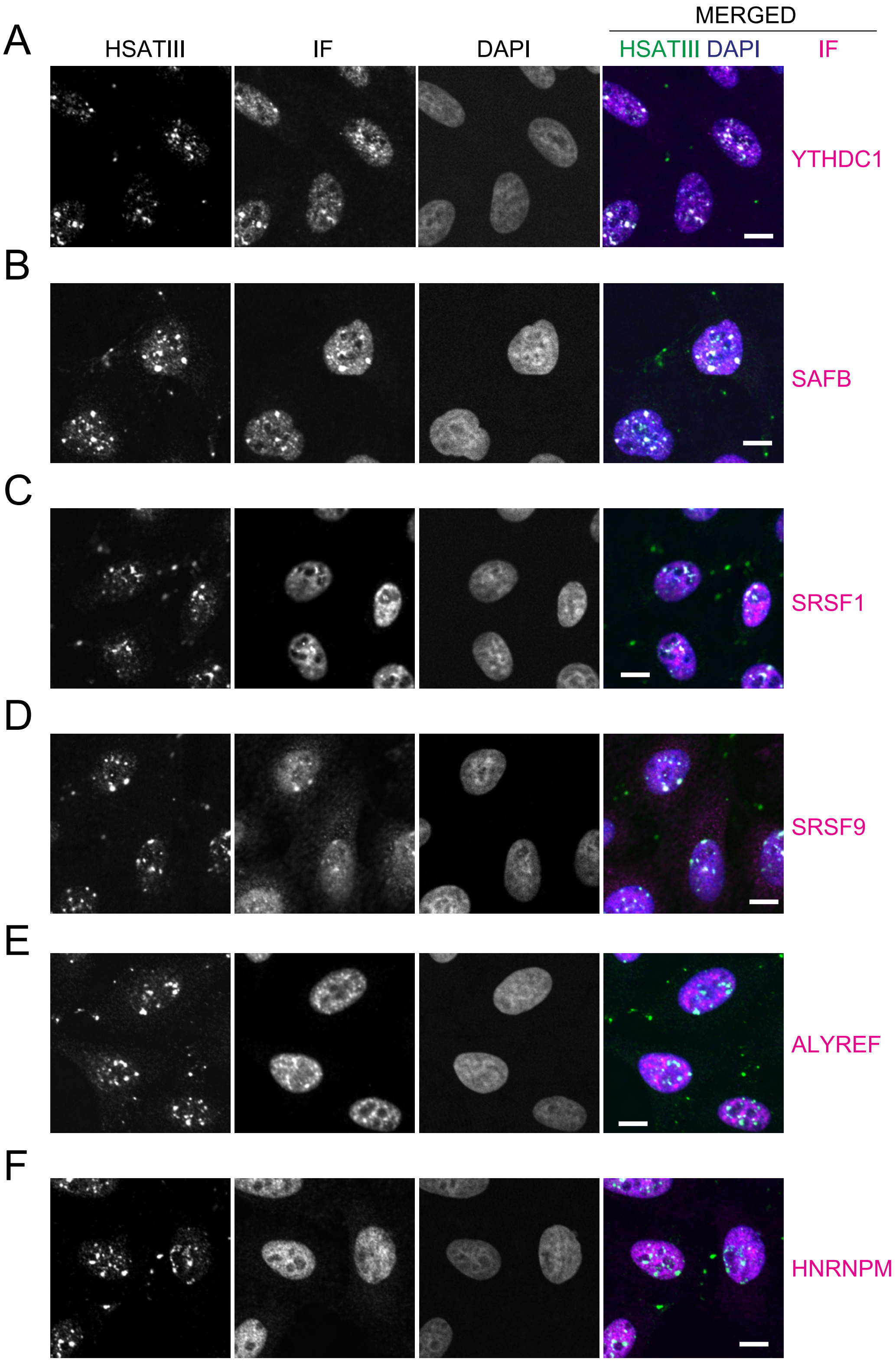

Supplemental Figure 3

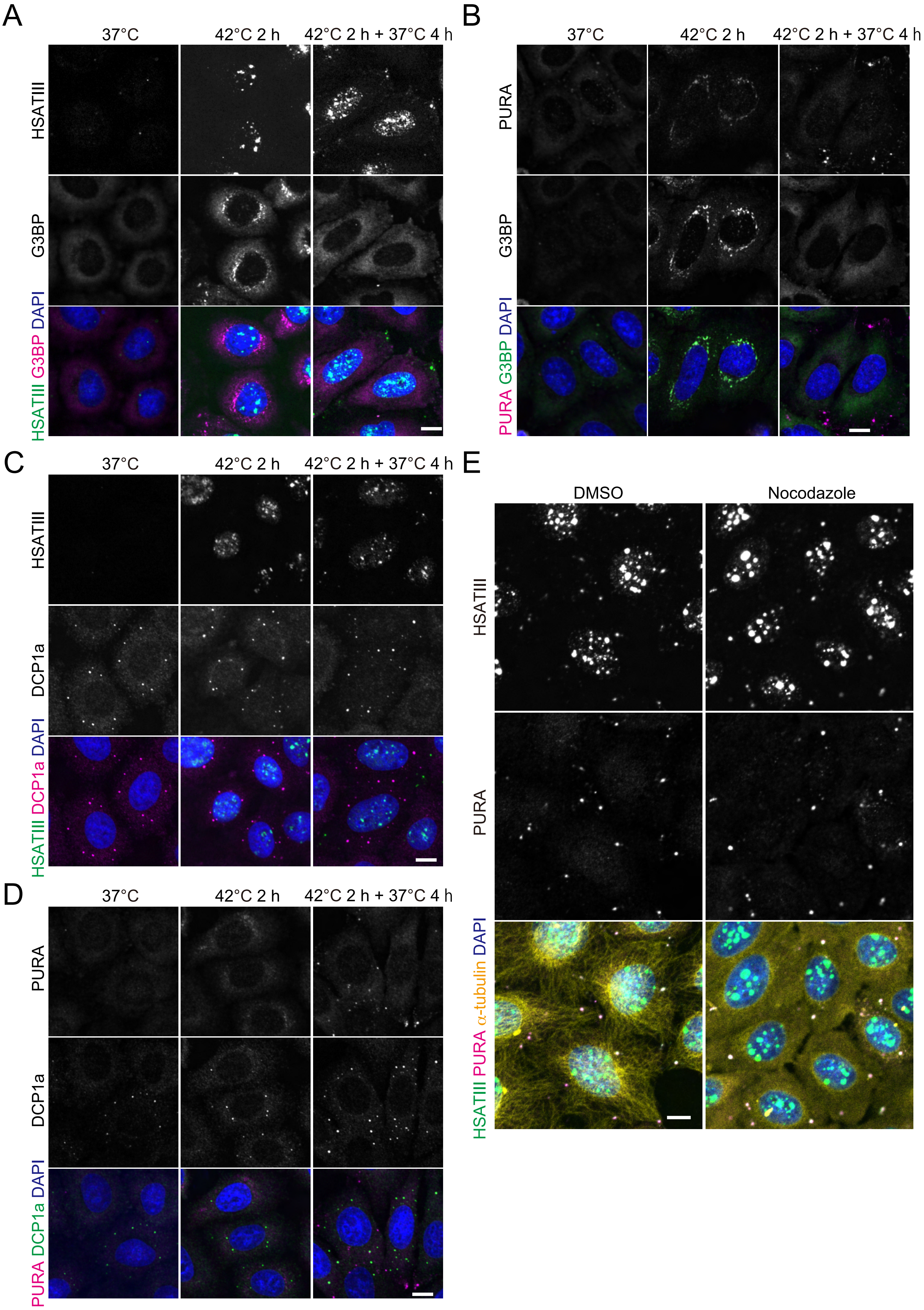

### Supplemental Figure 4

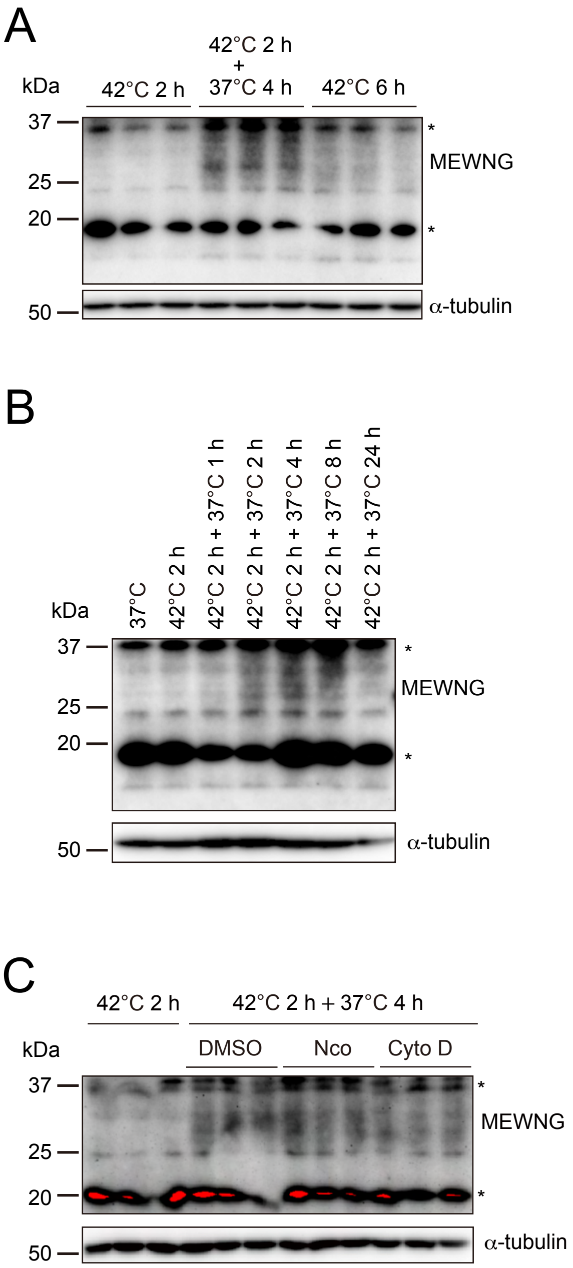

### Supplemental Figure 5

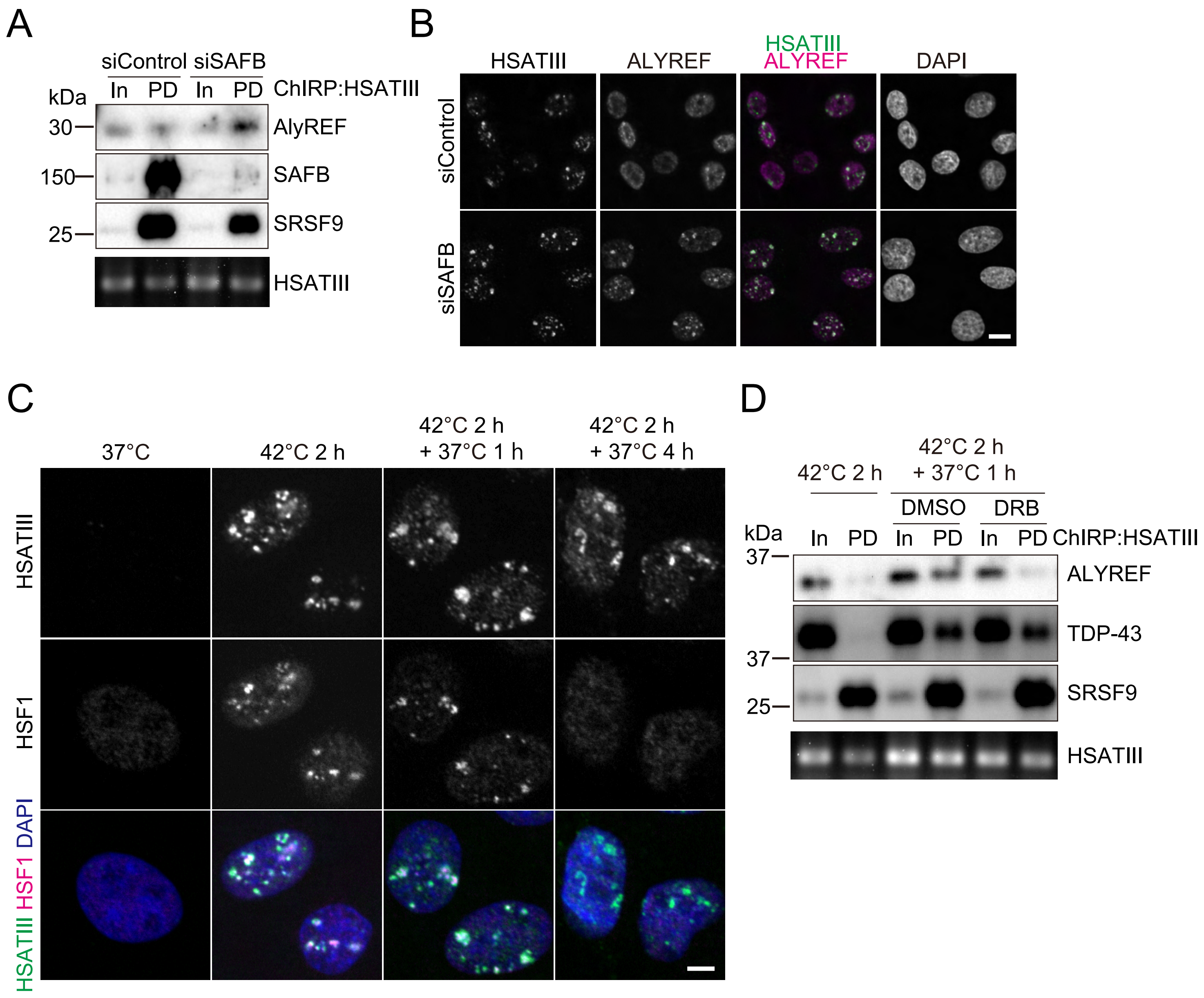

### Supplemental Figure 6

A

#8

MDYKDHDGDYKDHDIDYKDDDDTRAAASVMDICRIRPWNGRMEWNGMEWNGM
EWNGMEWMEWNGMEWNGMEWNGMEWKEWNGMEWNGMEWNGMEWNGM
EWNGMEWNGMEWNGMEWNGMEWNGMEWNGMEWNGMEWNGMEWNGME
WNGMEWNGMEWNGMEWNGMEWNGMEWNGMEWNGMEWNGMEWNGMEW
NGMEWNGMEWMEWNGMEWNGMEWNGMEWNGMEWNGMEWNGMEWNGM
EWNGMEWNGMEWNGMEWNGMEWNGMEWNGMEWNGMEWNGMEWNGMX
XXXXXX...

#32

MDYKDHDGDYKDHDIDYKDDDDTRAAASVMDICRIRPWNGWNGMEWNGINPN
GMEWNGMEWNGINPNGMEWNGMEWNGMESTRMEWNGMEWNGMESTRME
WNGMEWNGMESTRMEWNGMEWNGMESTRMEWNGMEWNGMESTRMEWNG
MEWNGMESTRMEWNGMEWNGMESTRMEWNGMEWNGMESTRMEWNGMEW
NGMESTRMEWNGMEWNGMESTRMEWNGMEWNGMESTRMEWNGMEWNGM
EWNQPEWNGMEWNGMEWNGINPNGMEWNGMEWNGINPNGMEWNGMESTR
MEWNGME

#12

MDYKDHDGDYKDHDIDYKDDDDTRAAASVMDICRIRPWNGINPNGMEWNGMES
TRWNGMEWNQPEWNGMEWNGINPNGMEWNGMESTRMEWNGMEWNQPEWN
GMEWNGMESTRMEWNGMEWNQPEWNGMEWNGINPNGMEWNGMESTRMEW
NGMEWNQPEWNGMEWNGINPNGMEWNGMESTRMEWNKGEFQHTGGRY

#59

MDYKDHDGDYKDHDIDYKDDDDTRAAASVMDICRIRPWNGMESTRMEWNGINP
NGMEWNQPEWNGMESTRMEWNGINPNGMEWNQPEWNGMESTRMEWNGIN
PNGMEWNQPEWNGMESTRMEWNGINPNGMEWNQPEWNGMESTRMEWNGI
NPNGMEWNQPEWNGMESTRMEWNGINPNGMEWNQPEWNGMESTRMEWNG
INPNGMEWNQPEWNGMESTRMEWNGINPNGMEWNQPEWNGMESTRMEWN
GINPNGMEWNQPEWNGME

B

#10

MDYKDHDGDYKDHDIDYKDDDDKLGTELGSNRINPNGMELNGMQWNGMEWN
GMEWYGIEWNGKKWNGMYSNQLEWNGMEWNCVEWNGMECKELNIINPNGM
EWNGMEWNGMESTRVEWNGMEWIGMECNEMEWNHSEWNGRECNRMEWK
GMESTRVDWKGMEWNGKEWNGINPSGMEWN

C

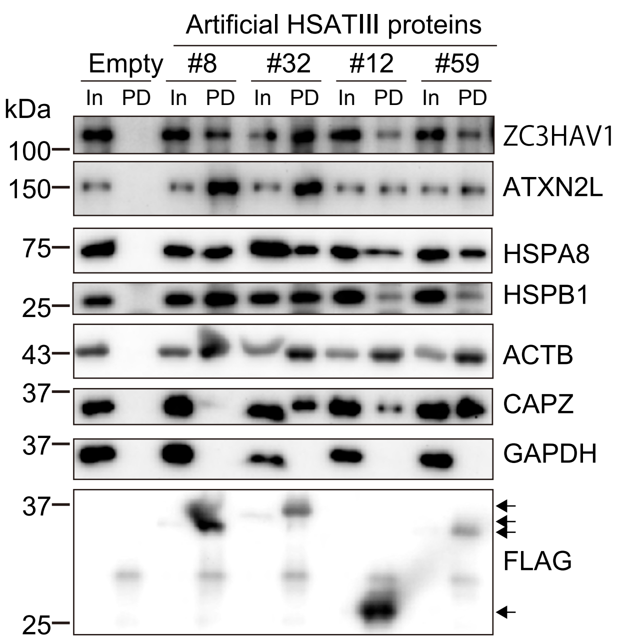


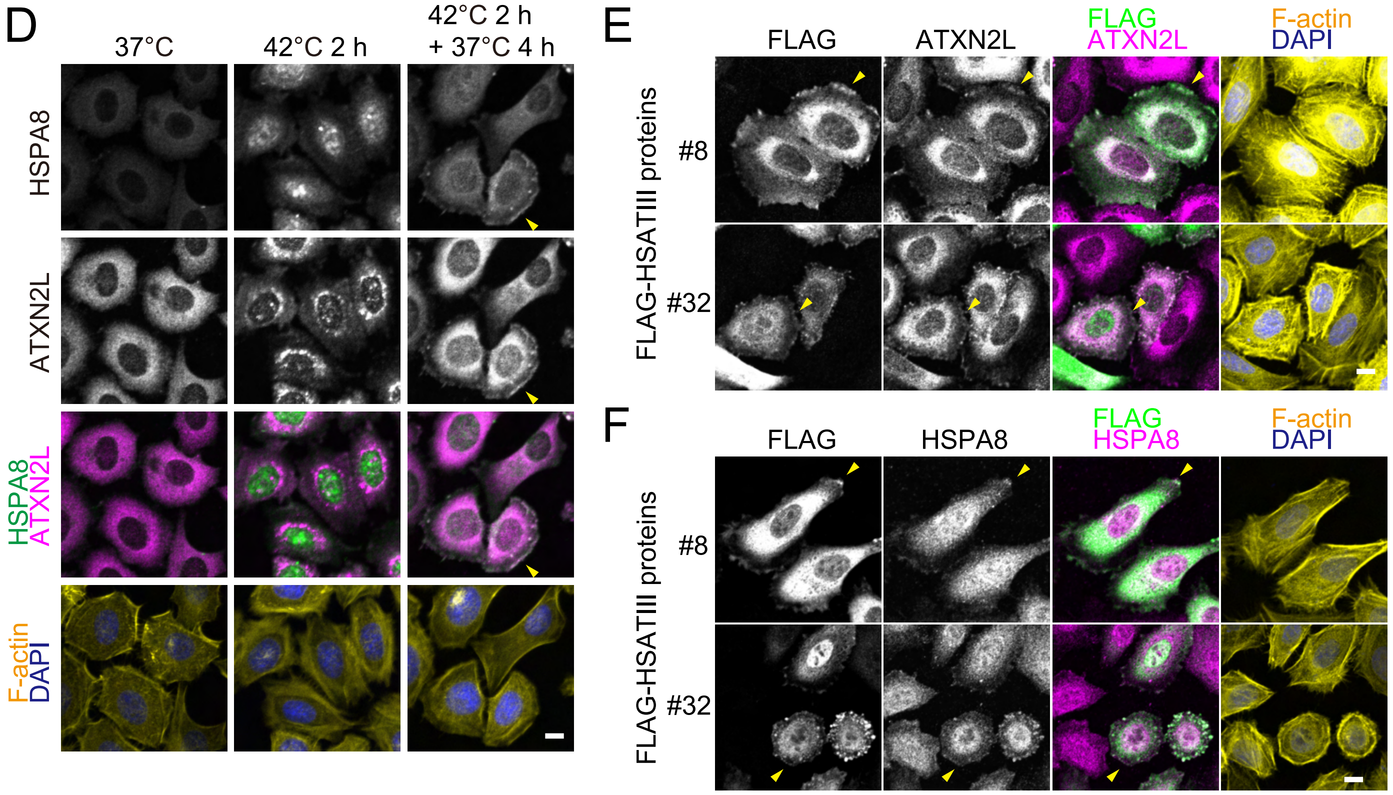

Supplemental Figure 7

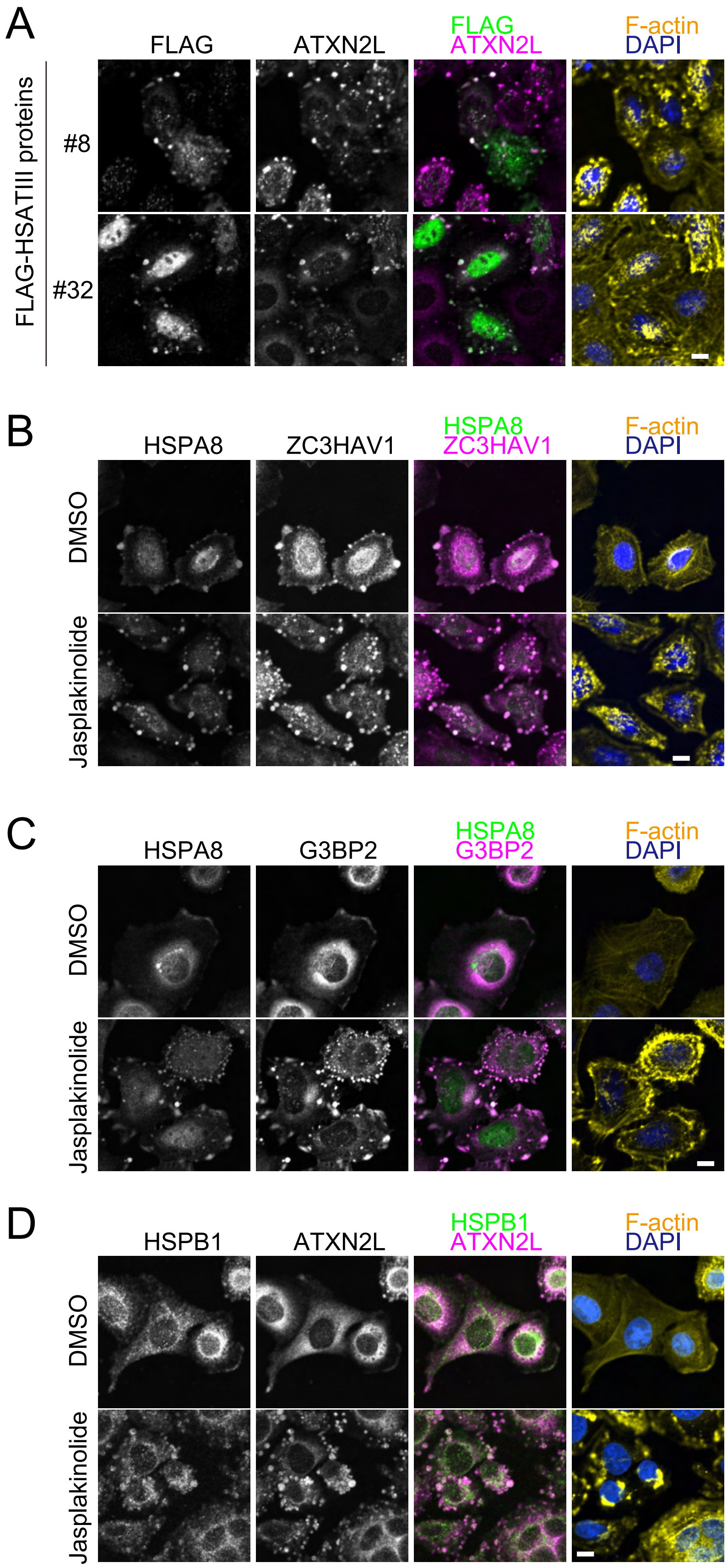

**Figure S1. Formation of cytoplasmic HSATIII RNA foci in post-stress recovery, related to Figure 1**

(A) Recovery phase specific formation of HSATIII RNA foci. HeLa cells were exposed to thermal stress as indicated and stained by HSATIII-FISH and IF using anti-PURA antibody. F-actin and nuclei were stained by phalloidin-iFluo$^{TM}$647 and DAPI, respectively. Scale bar: 10 μm

(B) HeLa cells were exposed to thermal stress (42 °C for 2 h followed by recovery at 37 °C for indicated hours). HSATIII lncRNAs were stained by FISH and IF as described above. Scale bar: 10 μm

**Figure S2. No localization of major nSB components in cytoplasmic HSATIII RNA foci related to Figure 2**

(A-F) HeLa cells (cultured at 42 °C for 2 h followed by recovery at 37 °C for 4 h) were stained by HSATIII-FISH and IF using a specific antibody against each protein. Scale bar: 10 μm

**Figure S3. Distribution of cytoplasmic HSATIII RNA foci and other cytoplasmic RNP foci, related to Figure 3**

(A-D) HeLa cells were cultured at 37°C (37°C) or at 42 °C for 2 h (42°C 2 h), followed by recovery at 37 °C for 4 h (42°C 2 h+ 37°C 4 h), and stained by HSATIII-FISH (A, C) and/or IF using a specific antibody against each protein. G3BP (A, B) and DCP1a (C, D) are used as markers of stress granules and p-bodies, respectively.

(E) Thermal stress-exposed HeLa cells (42°C 2 h+ 37°C 4 h) were treated with DMSO or Nocodazole (10 mg/ml) for 30 min and stained by HSATIII-FISH and IF using specific antibodies. Scale bar: 10 μm

**Figure S4. Western blotting analysis of HSATIII proteins, related to Figure 4**

(A) Recovery phase-dependent production of HSATIII proteins. HeLa cells were exposed to thermal stress as indicated and analyzed by western blot as described in Figure 4B.

(B) Time course analysis of HSATIII protein expression. HeLa cells exposed to thermal stress as indicated and analyzed by western blotting as described above.

(C) The effect of cytoskeleton disruption on HSATIII protein production. HeLa cells were treated as described in Figure 3D and analyzed by western blotting as described above.

**Figure S5. The mechanistic analyses of ALYREF localization to nSBs, related to Figure 5**

(A) The effect of SAFB KD on the ALYREF interaction with HSATIII RNAs. HeLa cells were transfected with siRNA against SAFB or control siRNA (10 nM), and cultured for 48 h, exposed to thermal stress (42 °C for 2 h, followed by recovery at 37 °C for 1 h), and analyzed by HSATIII-ChIRP. The ChIRP samples (pulldown, PD) were analyzed by western blot. HSATIII lncRNAs were detected by RT-PCR. Input (In) :1 % for WB, 100% for RT-PCR.

(B) The effect of SAFB KD on ALYREF localization to nSBs. HeLa cells prepared as described in (A) were stained by HSATIII-FISH and IF using ALYREF antibody. Nuclei were stained by DAPI. Scale bar: 10 μm

(C) HSF1 localization during thermal stress and recovery conditions. HeLa cells were exposed to thermal stress as indicated and stained by HSATIII-FISH and IF using HSF1 antibody. Nuclei were stained by DAPI. Scale bar: 10 μm

(D) The effect of transcription inhibitor on the ALYREF interaction with HSATIII RNAs. HeLa cells were exposed to thermal stress (42 °C for 2 h) and cultured at 37 °C for 1 h in the presence of DMSO or DRB (100 μM), and analyzed by HSATIII-ChIRP as described in (A). Input (In) :1 % for WB, 100% for RT-PCR.

**Figure S6. Interaction and colocalization of HSATIII proteins with the binding proteins, related to Figure 6**

(A, B) The amino acid sequences of artificially-designed HSATIII (adHSATIII) proteins (A) and exogenous HSATIII-derived protein (clone #10, B). Also see the DNA sequences in Table S2. FLAG-tag, MEWNG region and low frequent repeat-derived amino acids are indicated in green, magenta and cyan, respectively. Underlines indicate vector sequence-derived regions. “XXXX…” in adHSATIII #8 indicates the sequence-undetermined region.

(C) IP-western blot validation of HSATIII protein-binding proteins. IP was performed using anti-FLAG antibody from HeLa cells (42 °C 2 h + 37 °C 4 h) expressing FLAG-tagged artificial HSATIII proteins in (A). Western blot analysis was performed using indicated antibodies. GAPDH was used as negative control. In (Input 1%), PD (pulldown). An asterisk shows IgG light chain. Arrows indicates the positions of FLAG-tagged HSATIII proteins

(D) Subcellular localization of HSPA8 and ATXN2L in thermal stress response. HeLa cells at indicated conditions were stained by IF using specific antibodies. F-actin and nucleus were stained by phalloidin-iFlour$^{TM}$647 and DAPI, respectively. Scale bar: 10 $\mu$m

(E, F) Colocalization of adHSATIII proteins (adHSATIII #8 and #32) with ATXN2L (E) and HSPA8 (F). The experiments were performed as described in Figure 6G and 6H. Scale bar: 10 $\mu$m

**Figure S7. Actin polymerization-dependent aggregation of HSATIII protein complexes, related to Figure 7**

(A) Actin polymerization-dependent aggregation of HSATIII protein complexes. HeLa cells expressing adHSATIII proteins (adHSATIII #8 and #32) were prepared and stained same as Figure7A. HeLa cells were treated with jasplakinolide (100 nM, 30 min) before fixation. Scale bar: 10 μm

(B-D) Actin polymerization dependent co-aggregation of HSATIII protein-binding proteins. HeLa cells (42 °C 2 h + 37 °C 4 h) were treated with DMSO or jasplakinolide (100 nM, 30 min) before fixation and stained by IF. F-actin and nucleus were stained by phalloidin-iFlour™647 and DAPI, respectively. Scale bar: 10 μm
