## Supplemental Table S2 for "Cytoplasmic roles of HSATIII RNAs in RNA granule assembly and production of actin cytoskeleton-associated repeat-containing proteins"

### Table S2. DNA sequences of endogenous and artificially designed HSATIII clones

| Clone | Sequence | Note |
|---|---|---|
| #10 | AATAGAATCAACCCGAATGGAATGGAATTGAACGGAATGCAATGGAATGGAATGGA<br>ATGGAATGGAATGGAATGGTACGGAATAGAATGGAATGGAAAGAAATGGAATGGAA<br>TGTATTCAAACCAATTGGAATGGAATGGAATGGAATGGAATTGTGTGGAATGGAAT<br>GGAATGGAGTGTAAAGAATTGAATATAATCAACCCGAATGGAATGGAATGGAATGG<br>AATGGAATGGAATGGAATGGAATCAACTCGAGTGGAATGGAATGGTATGGAATGGA<br>TTGGAATGGAATGCAATGAAATGGAATGGAATCATTCCGAGTGGAATGGAAGGGAA<br>TGTAATCGAATGGAATGGAAAGGAATGGAATCAACCCGAGTGGATTGGAAGGGAA<br>TGGAATGGAATGGAAAGGAATGGAATGGAATCAACCCGAGTGGAATGGAATGGAA<br>TTGAATGGAATATAATCA | |
| #8 | GGAATGGAAGAATGGAATGGAATGGAATGGAATGGAATGGAATGGAATGGAATGG<br>AATGGAATGGATGGAATGGAATGGAATGGAATGGAATGGAATGGAATGGAATGGAA<br>TGGAATGGAAGGAATGGAATGGAATGGAATGGAATGGAATGGAATGGAATGGAAT<br>GGAATGGAATGGAATGGAATGGAATGGAATGGAATGGAATGGAATGGAATGGAATG<br>GAATGGAATGGAATGGAATGGAATGGAATGGAATGGAATGGAATGGAATGGAATGG<br>AATGGAATGGAATGGAATGGAATGGAATGGAATGGAATGGAATGGAATGGAATGGA<br>ATGGAATGGAATGGAATGGAATGGAATGGAATGGAATGGAATGGAATGGAATGGAA<br>TGGAATGGAATGGAATGGAATGGAATGGAATGGAATGGAATGGAATGGAATGGAAT<br>GGAATGGAATGGAATGGAATGGAATGGAATGGAATGGAATGGAATGGAATGGAATG<br>GAATGGAATGGATGGAATGGAATGGAATGGAATGGAATGGAATGGAATGGAATGGA<br>ATGGAATGGAATGGAATGGAATGGAATGGAATGGAATGGAATGGAATGGAATGGAA<br>TGGAATGGAATGGAATGGAATGGAATGGAATGGAATGGAATGGAATGGAATGGAAT<br>GGAATGGAATGGAATGGAATGGAATGGAATGGAATGGAATGGAATGGAATGGAATG<br>GAATGGAATGGAATGGAATGGAATGGAATGGATGG | The length is putative, because the precise number of repeat units in the middle region could not be determined. |
| #32 | GGAATGGATGGAATGGAATGGAATGGAATGGAATCAACCCGAATGGAATGGAATG<br>GAATGGAATGGAATGGAATGGAATCAACCCGAATGGAATGGAATGGAATGGAATGG<br>AATGGAATGGAATGGAATCAACCCGAATGGAATGGAATGGAATGGAATGGAATGGA<br>ATGGAATCAACCCGAATGGAATGGAATGGAATGGAATGGAATGGAATGGAATCAAC<br>CCGAATGGAATGGAATGGAATGGAATGGAATGGAATGGAATCAACCCGAATGGAAT<br>GGAATGGAATGGAATGGAATGGAATGGAATCAACCCGAATGGAATGGAATGGAATG<br>GAATGGAATGGAATGGAATCAACCCGAATGGAATGGAATGGAATGGAATGGAATGG<br>AATGGAATCAACCCGAATGGAATGGAATGGAATGGAATGGAATGGAATGGAATCAA<br>CCCGAATGGAATGGAATGGAATGGAATGGAATGGAATGGAATCAACCCGAATGGA<br>ATGGAATGGAATGGAATGGAATGGAATGGAATCAACCCGAATGGAATGGAATGGAA<br>TGGAATGGAATGGAATGGAATCAACCCGAATGGAATGGAATGGAATGGAATGGAAT | |

| | |
|---|---|
| | GGAATGGAATGGAATCAACCCGAATGGAATGGAATGGAATGGAATGGAATGGAATG<br>GAATGGAATCAACCCGAATGGAATGGAATGGAATGGAATGGAATGGAATGGAATCA<br>ACCCGAATGGAATGGAATGGAATGGAATGGAATCAACCCGAATGGAATGGAATGG<br>AATGGAATGGAATGGAATGGAAT |
| #12 | GGAATGGAATCAACCCGAATGGAATGGAATGGAATGGAATGGAATCAACCCGATG<br>GAATGGAATGGAATGGAATCAACCCGAATGGAATGGAATGGAATGGAATGGAATCA<br>ACCCGAATGGAATGGAATGGAATGGAATGGAATCAACCCGAATGGAATGGAATGG<br>AATGGAATGGAATCAACCCGAATGGAATGGAATGGAATGGAATGGAATGGAATCAA<br>CCCGAATGGAATGGAATGGAATGGAATGGAATCAACCCGAATGGAATGGAATGGA<br>ATGGAATGGAATCAACCCGAATGGAATGGAATGGAATGGAATGGAATCAACCCGAA<br>TGGAATGGAATGGAATGGAATGGAATCAACCCGAATGGAATGGAATGGAATGGAAT<br>GGAATCAACCCGAATGGAATGGAATGGAATGGAATGGAATCAACCCGAATGGAATG<br>GAAT |
| #59 | GGAATGGAATGGAATCAACCCGAATGGAATGGAATGGAATCAACCCGAATGGAATG<br>GAATGGAATCAACCCGAATGGAATGGAATGGAATCAACCCGAATGGAATGGAATGG<br>AATCAACCCGAATGGAATGGAATGGAATCAACCCGAATGGAATGGAATGGAATCAA<br>CCCGAATGGAATGGAATGGAATCAACCCGAATGGAATGGAATGGAATCAACCCGA<br>ATGGAATGGAATGGAATCAACCCGAATGGAATGGAATGGAATCAACCCGAATGGAA<br>TGGAATGGAATCAACCCGAATGGAATGGAATGGAATCAACCCGAATGGAATGGAAT<br>GGAATCAACCCGAATGGAATGGAATGGAATCAACCCGAATGGAATGGAATGGAATC<br>AACCCGAATGGAATGGAATGGAATCAACCCGAATGGAATGGAATGGAATCAACCC<br>GAATGGAATGGAATGGAATCAACCCGAATGGAATGGAATGGAATCAACCCGAATGG<br>AATGGAATGGAATCAACCCGAATGGAATGGAATGGAATCAACCCGAATGGAATGGA<br>ATGGAATCAACCCGAATGGAATGGAATGGAATCAACCCGAATGGAATGGAATGGAA<br>TCAACCCGAATGGAATGGAATGGAATCAACCCGAATGGAATGGAATGGAATCAACC<br>CGAATGGAATGGAATGGAAT |
