## Supplemental Table S3 for "Cytoplasmic roles of HSATIII RNAs in RNA granule assembly and production of actin cytoskeleton-associated repeat-containing proteins"

### Table S3. The list of primers used in this paper, related to STAR Methods

| | | |
|---|---|---|
| **PCR primers for RT-PCR (5'→3')** | | |
| HSATIII | TATGAATTCAATCAACCCGAGTGCAATCGAA | TATGGATCCTTCCATTCCATTGCTGTACTCG |
| GAS5 | ATGCAGTGTGGCTCTGGATAGC | AAGCTGGTCCAGGCAAGTTGGAC |
| 18S ribosomal RNA | ATTAACAACGAAAGTCGGAGGT | TTTAAGTTTCAGCTTTGCAACCATACT |
| **PCR primers for vector construction (5'→3')** | | |
| HSATIII fragment #10 | GAGCTCGGATCCAATAGAATCAACCCGAATGG | TCTAGAGCGGCCGCTGATTATATTCCATTCAATTCC |
| TurboID vector | TACCGG TCTAGAATGGCTAGCAAAGAC | TCAGATCTCGAGTCTGCAGCTTTTCGGCAGACCGCAG |
| **Primers for repeat extension PCR (5'→3')** | | |
| (GGAAT)$_7$CAACCCGAAT(GGAAT)$_7$ | GGAATGGAATGGAATGGAATGGAATGGAATGGAATCAACCCGAATGGAATGGAATGGAATGGAATGGAATGGAATGGAAT | ATTCCATTCCATTCCATTCCATTCCATTCCATTCCATTCGGGTTGATTCCATTCCATTCCATTCCATTCCATTCCATTCC |
| (GGAAT)$_5$CAACCCGAAT(GGAAT)$_5$ | GGAATGGAATCAACCCGAATGGAATGGAATGGAATGGAATGGAATCAACCCGAATGGAATGGAATGGAATGGAATGGAATCAACCCGAATGGAATGGAAT | ATTCCATTCCATTCGGGTTGATTCCATTCCATTCCATTCCATTCCATTCGGGTTGATTCCATTCCATTCCATTCCATTCCATTCGGGTTGATTCCATTCC |
| (GGAAT)$_3$CAACCCGAAT(GGAAT)$_3$ | GGAATGGAATGGAATCAACCCGAATGGAATGGAATGGAATCAACCCGAATGGAATGGAATGGAATCAACCCGAATGGAATGGAATGGAATCAACCCGAATGGAATGGAATGGAAT | ATTCGGGTTGATTCCATTCCATTCCATTCGGGTTGATTCCATTCCATTCCATTCGGGTTGATTCCATTCCATTCCATTCGGGTTGATTCCATTCCATTCCATTCCATTCCATTCC |
| (GGAAT)$_{20}$ | GGAATGGAATGGAATGGAATGGAATGGAATGGAATGGAATGGAATGGAATGGAATGGAATGGAATGGAATGGAATGGAATGGAATGGAATGGAATGGAAT | ATTCCATTCCATTCCATTCCATTCCATTCCATTCCATTCCATTCCATTCCATTCCATTCCATTCCATTCCATTCCATTCCATTCCATTCCATTCCATTCC |
